# DFCP1 is a Regulator of ATGL-mediated Lipid Droplet Lipolysis

**DOI:** 10.1101/2024.02.14.580382

**Authors:** V.A. Ismail, M.N. Harvey, J. Castillo-Badillo, T. Naismith, D.J. Kast

## Abstract

Lipid droplets (LDs) are transient lipid storage organelles that can be readily tapped to resupply cells with energy or lipid building blocks, and therefore play a central role in cellular metabolism. Double FYVE Domain Containing Protein 1 (DFCP1/ZFYV1) has emerged as a key regulator of LD metabolism, where the nucleotide-dependent accumulation of DFCP1 on LDs influences their size, number, and dynamics. Here we show that DFCP1 regulates lipid metabolism by directly modulating the activity of Adipose Triglyceride Lipase (ATGL/PNPLA2), the rate-limiting lipase driving the catabolism of LDs. We show through pharmacological inhibition of key enzymes associated with LD metabolism that DFCP1 specifically regulates lipolysis and, to a lesser extent, lipophagy. Consistent with this observation, DFCP1 interacts with and recruits ATGL to LDs in starved cells, irrespective of known regulatory factors of ATGL. We further establish that this interaction prevents dynamic disassociation of ATGL from LDs and thereby impedes the rate of LD lipolysis. Collectively, our findings indicate that DFCP1 primes ATGL on LDs to promote rapid LD catabolism.

## Introduction

Lipid droplets (LDs) are transient lipid storage depots that provide lipids for the repair and biogenesis of membranous organelles, while also serving as a reservoir of energy for the cell. The breakdown of triacylglycerides (TAGs) from LDs into fatty acids (FAs) during nutrient stress plays a critical role in cell survival, and failure to do so contributes to the pathology of a host of metabolic diseases including lipodystrophies, obesity, insulin resistance, diabetes, NAFLD, and atherosclerosis (1). Thus, it is important for the cell to maintain tight control over LD metabolism via many regulatory proteins across key steps of LD growth and degradation.

The process of LD metabolism is incredibly dynamic and involves the packaging of TAGs into LDs (LD biogenesis) and the stimulated breakdown of the TAGs stored inside LDs to release FAs (LD catabolism) that are used by mitochondria. The latter stage is thought to be mediated by two sequential and intertwined mechanisms involving breakdown of large LDs by LD-associated lipases into small LDs via the lipolytic pathway (lipolysis), followed by efficient clearance by the autophagy-lysosomal pathway, in a distinct process known as lipophagy (2, 3). It is believed that the lipophagy pathway can only break down smaller LDs due to the limited size of the phagophore, and thus, cells that form larger LDs tend to favor lipolysis to mobilize FAs (4). It should be noted that a surplus of FAs can also be repackaged into LDs, which is important for protecting cells from the toxic accumulation of FAs in the cytosol.

During lipolysis, TAGs are sequentially hydrolyzed into diacylglycerol (DAG), monoacylglycerol (MAG), and glycerol, releasing a FA at each of these steps. The first step of this process - hydrolysis of TAG into DAG - is slow and requires the lipase adipose triglyceride lipase (ATGL). However, the process can be markedly enhanced through a combination of AMPK and/or PKA-dependent phosphorylation of ATGL (5) (6), as well as through interaction of ATGL with its coactivator 1-acylglycerol-3-phosphate o-acyltransferase (ABHD5), more commonly known as CGI-58 (7). Aside from phosphorylation and regulation via a few key regulatory proteins, very little is known about the molecular factors that modulate ATGL’s lipolytic activity.

The autophagy-associated protein, DFCP1, has recently gained a lot of attention as a novel regulator of LD metabolism. Both endogenous (8) and overexpressed (8–10) DFCP1 has been shown to accumulate on LDs in cells, and the expression of DFCP1 correlates with increased LD size (9), whereas knockdown of DFCP1 was shown to do the opposite – it significantly decreased LD size while concomitantly increasing the number of LDs. This accumulation of LDs was also shown to be dependent on a novel bifunctional NTPase domain housed within DFCP1 that hydrolyzes both ATP and GTP (8). Additionally, it was proposed that DFCP1 is recruited along with Rab18 and Seipin to LDs through an interaction with ZW10 (Centromere/kinetochore protein zw10 homolog) (11), a central component of the NRZ ER-LD contact site complex. However, it is unknown if DFCP1’s functions on LDs requires the NRZ complex, or if DFCP1 can act as an autonomous tether between LDs and the ER.

The specific mechanism by which DFCP1 regulates LD size is unclear. It has been proposed that DFCP1 participates in the biogenesis of LDs shortly after the import of exogenous FAs. The addition of FAs has been shown to promote an autophagic response (12, 13), which leads to an increase in PI3P-rich autophagosome precursor sites in the ER. Consequently, DFCP1 accumulates at these sites due to its intrinsic ability to bind PI3P. While DFCP1 was shown to not influence autophagosome formation (8, 14), it may however assist in the biogenesis of LDs at these same sites (15). Recently, DFCP1 has been shown to promote TAG storage in LDs by impairing LD turnover and FA metabolism (8). However, during times of nutrient stress, DFCP1 partially translocates away from LDs to sites of autophagosome formation on the ER in a manner that depends on its NTPase activity, which promotes LD turnover and an increase of FA metabolism (8). This suggests that when DFCP1 is associated with LDs, it directly impairs lipolysis, while DFCP1 at PI3P-rich regions slightly impedes the lipophagic breakdown of LDs. However, the mechanism by which DFCP1 regulates lipolysis remains unknown.

Here we show that DFCP1 is a regulator of the rate-limiting lipolysis enzyme, ATGL. Specifically, we show that ablation of DFCP1 profoundly influences LD turnover in cells that were pharmacologically inhibited in their ability to perform LD biogenesis and lipophagy. Moreover, we found that DFCP1 directly interacts with ATGL via the N-terminus of ATGL and the C-terminus of DFCP1, and the interaction is important for both the recruitment and sequestration ATGL to the LD surface. Conversely, ablation of DFCP1 markedly enhances ATGL dynamics on LDs and leads to a marked increase in the in the lipolysis of LDs. Overall, we demonstrate that DFCP1 primes lipolysis by anchoring and suppressing ATGL on the surface of LDs. Upon starvation and/or nucleotide hydrolysis, DFCP1 releases this repression and allows for ATGL to begin lipolysis.

## Results

### DFCP1 Inhibits LD Catabolism

DFCP1 has been implicated in regulating LD biogenesis (9, 15) and catabolism (8) where the knockdown (KD) of DFCP1 is associated with smaller, more numerous LDs. Interestingly, we noted that in U2OS cells, stimulated with 200 μM oleic acid (OA), LD diameters increase as DFCP1 levels increase in the cell **(Figure 1A and 1B)**, particularly when cells are starved of amino acids by replacing the growth medium with EBSS for 4 or more hours (hereafter referred to as “starved”). This effect is exacerbated when DFCP1 is overexpressed and suggests that DFCP1 inhibits degradation of large LDs during starvation, a condition that drives the catabolism of LDs. However, given that LD biogenesis and the two canonical catabolic pathways, lipolysis and lipophagy occur simultaneously, it is also possible that DFCP1 promotes biogenesis and/or accelerates maturation of LDs in cells. Therefore, it is difficult to determine DFCP1’s precise role in LD metabolism without systematically examining the impact of DFCP1 on biogenesis, lipolysis and lipophagy.

**Figure 1:**
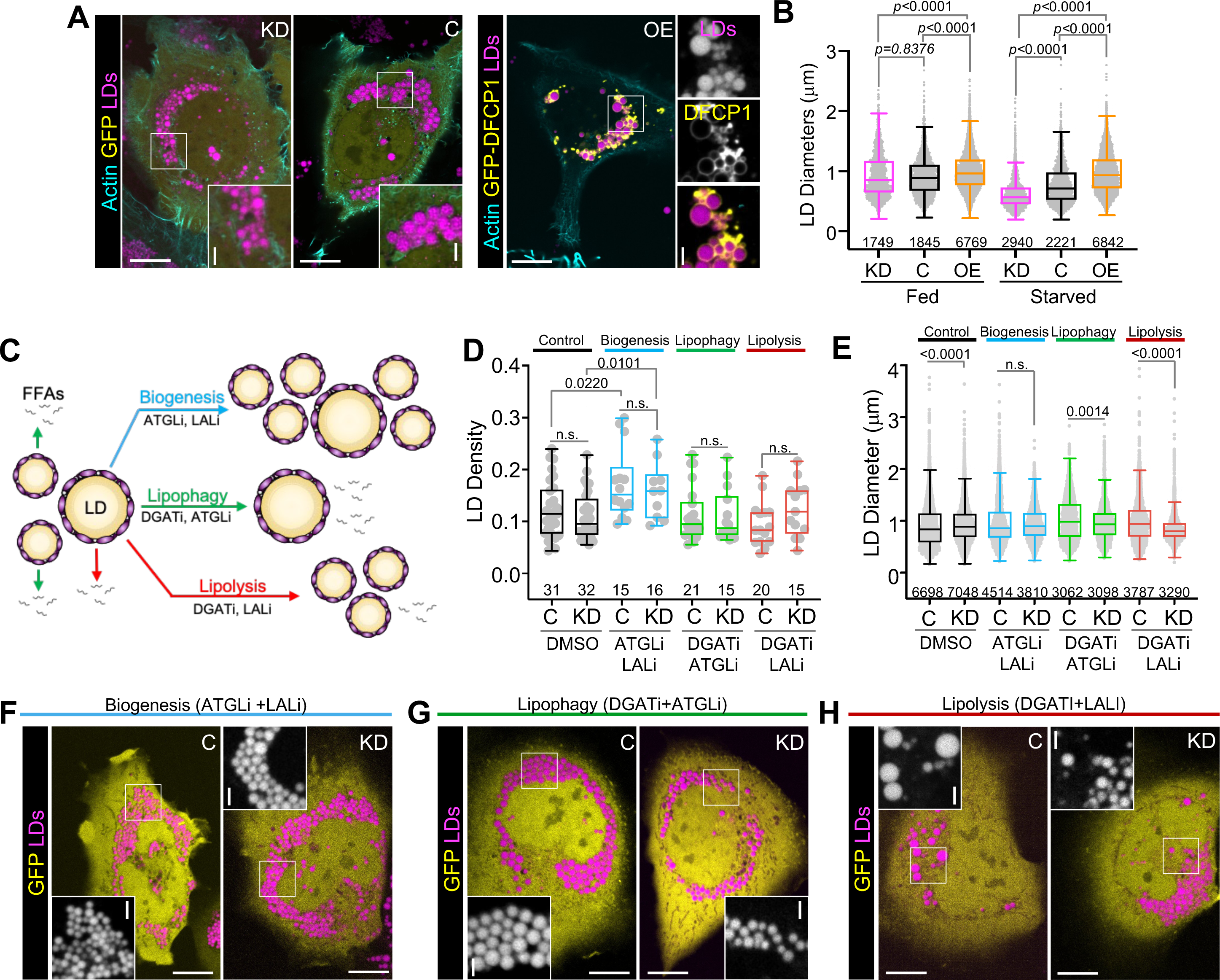
DFCP1 Inhibits LD Catabolism. **(A)** Live cell confocal images of U2OS cells treated with DFCP1 siRNA (KD) or non-targeting siRNA treatment (control or C), expressing GFP-DFCP1 or GFP (yellow), and BFP-LifeAct (cyan). Cells were treated with 200 µM oleic acid (OA) for 20 h to stimulate LD formation. Prior to imaging, cells were incubated for 4 h in starvation media (EBSS) and then stained with LipidTOX Deep Red for 30 min. **(B)** LD diameter distributions of LDs from cells treated in **A**. **(C)** Diagram of LD growth and degradation pathways, including the combinations of small molecule inhibitors used to isolate specific pathways in D-H. **(D)** Number of LDs/cell area (density) quantified from images of control and DFCP1 KD cells expressing GFP that were treated with 200 µM OA for 20 h and starved (EBSS) for 18 h along with DMSO (vehicle) or the indicated inhibitors. **(E)** LD diameter quantified from images of cells that were treated in **D**. **(F-G)** Representative images of cells quantified in **D** and **E**. The scale bars in whole-cell and inset images represent 10 and 2 µm, respectively. The statistical significance of the measurements in **B, D and E** was determined using the Mann–Whitney U-test on the indicated number of observations from two independent transfections. Exact *p*-values are reported with exception to *p*>0.05, which is not considered significant (n.s.).

To address this question, we examined the effect of knocking down DFCP1 with siRNA on the number and size of LDs in response to treating cells with a combination of well-established small molecule inhibitors for diglyceride acyltransferase 1 (diacylglycerol acetyl transferase 1 or DGAT1), adipose triglyceride lipase or ATGL and/or lysosomal acid lipase or LAL, which are key enzymes involved in LD biogenesis, lipolysis or lipophagy, respectively (16–18). By multiplexing these inhibitors, it is possible to specifically block biogenesis, lipophagy, or lipolysis (**Figure 1C**). Indeed, control cells (cells treated with a non-targeting siRNA) that were starved for 4 h after a 24 h treatment of OA and a combination of the ATGL inhibitor (ATGLi) and the inhibitor for LAL (LALi), had a significantly higher density of LDs (number of LDs normalized by the volume of the cell) but the diameters of these LDs were unchanged from DMSO (vehicle) treated cells (**Figure 1D-1F)**. This indicates that under these conditions, cells are able to generate LDs, but the mechanism to reduce their size or remove them has been stalled. By contrast, starving cells treated with OA and a combination of the DGATi and either ATGLi (**Figure 1D, 1E and 1G**) or LALi (**Figure 1D, 1E, and 1H**) led to a small decrease in LD density, but a marked increase in diameters of these LDs when compared to DMSO treated cells, which is consistent with blocking biogenesis (reduced number) and one of the catabolic processes (increased size).

Having established that we could selectively monitor only one active pathway of LD metabolism at a time, we then examined the impact of these combinations of inhibitors on starved OA treated cells where DFCP1 was knocked down. In starved KD cells treated with a combination of OA, ATGLi and LALi, there was not a significant change in LD density or diameter when compared to control cells treated in the same way **(Figure 1D-F).** This was quite surprising since DFCP1 has been previously implicated in LD biogenesis (15). Thus, we focused our attention on the catabolic pathways. When we investigated lipophagy by blocking biogenesis in conjunction with lipolysis, using the DGATi and ATGLi inhibitors (**Figure 1G**), we did not observe a difference in LD density (**Figure 1D**), when compared to control cells that were treated with the same conditions, but we did observe a small but significant decrease in the distribution of LD diameters (**Figure 1E**). This is consistent with our previous observations on lipophagy, where we observed that DFCP1 KD reduced the number of LDs associated with autophagosomes, which results in accumulation of smaller LDs (8). Finally, when we examined the impact of DFCP1 KD on lipolysis by blocking both biogenesis and lipophagy with DGATi and LALi, respectively, we found that DFCP1 KD cells (**Figure 1H**) exhibited a higher density of LDs, that were significantly smaller than those in control treated cells (**Figure 1D and 1E**). This latter observation suggests that DFCP1 may function early in the pathway of LD catabolism, where it could protect LDs from lipolysis, which would in turn impair lipophagy. For this reason, we decided to focus our attention on defining the role of DFCP1 in lipolysis.

### DFCP1 Controls ATGL Localization in Cells

LD lipolysis involves the sequential hydrolysis of TAGs into FFAs and glycerol derivatives through the sequential activities of ATGL, hormone sensitive lipase (HSL) and monoglyceride lipase (MGL). Among these enzymes, ATGL is known to be rate-limiting, and thus factors that regulate ATGL often have a profound effect on lipid storage. ATGL is of particular interest here because DFCP1 was initially discovered to be associated with LDs using proximity ligation approaches in combination with mass spectrometry (19). That study also reported that DFCP1 was in proximity of ATGL but not PLPN2, an LD coat protein, which may indicate that DFCP1 is not simply an LD-associated protein but may function with ATGL in some capacity. To gain insight into this possibility, we examined the localization of ATGL to LDs in function of nutrient conditions and DFCP1 expression. For this purpose we ablated DFCP1 in U2OS cells using CRISPR/Cas9 (8), which did not significantly impact the viability (**Supplemental Figure 1A**) or the number of cells entering early (annexin V) or late (propidium iodide) apoptosis (**Supplemental Figure 1B**) when exposed to exogenous oleic (OA) or palmitic acid (PA) under fed or starved conditions. Using these cells, we found that ablation of DFCP1 had little effect on the localization of ATGL to LDs in fed cells that were treated with 200 μM OA for 20 h, but, instead, significantly diminished the colocalization of ATGL with LDs in starved cells (**Figure 2A and 2B**). Conversely, the overexpression of BFP-DFCP1 markedly increased the localization of GFP-ATGL to LDs in starved cells, when compared to cells overexpressing BFP alone (**Figure 2C and 2D**). This DFCP1-dependent recruitment of ATGL to LDs was also dependent on the nutritional status of the cell, since we saw more colocalization of GFP-ATGL with BFP-DFCP1^WT^ when cells were starved when compared to those that were fed (**Figure 2E**). Coincidently, we also observed an increase in localization of BFP-DFCP1^WT^ to LDs in starved cells, when compared to fed cells expressing GFP-ATGL (**Supplemental Figure 1C**). This is particularly surprising, because we have previously shown that starvation partially relocalizes DFCP1 from LDs to sites of autophagosome biogenesis (8), and thus suggests that increased expression of either protein may mutually reinforce their localization to LDs during starvation.

**Figure 2:**
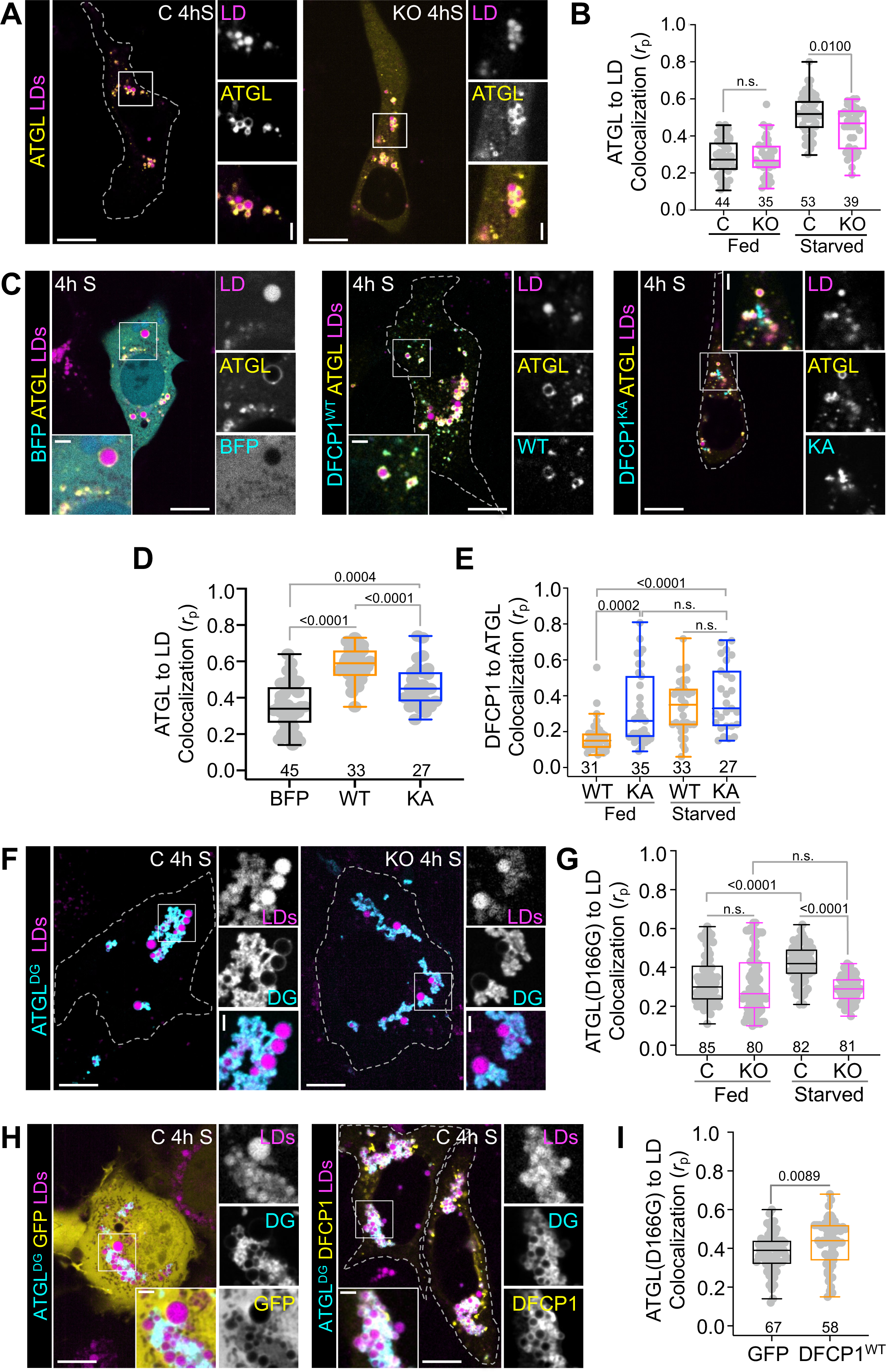
DFCP1 Controls ATGL Localization in Cells. **(A)** Representative images of U2OS control (C) and DFCP1 CRISPR KO cells (KO) expressing GFP-ATGL and treated with 200 µM OA for 20 h and then fed (basal growth media) or starved (EBSS) for 4 h and treated with LipidTOX Deep Red for 30 min prior to imaging. **(B)** Colocalization (Pearson’s correlation coefficient, *rp*) of GFP-ATGL with LDs in control (black) and KO (magenta) cells described in **A**. **(C)** Representative images of U2OS cells expressing GFP-ATGL and either BFP, BFP-DFCP1^WT^ or BFP-DFCP1^KA^. Cells were treated with OA and starved as in A. **(D)** Colocalization (Pearson’s correlation coefficient) of GFP-ATGL with LDs in cells described in **C** that were transfected with GFP-ATGL and BFP (black), BFP-DFCP1^WT^ (WT, orange) or BFP-DFCP1^KA^ (KA, blue). **(E)** Colocalization (Pearson’s correlation coefficient, *rp*) of GFP-ATGL with either BFP-DFCP1^WT^ (WT, orange) or BFP-DFCP1^KA^ (KA, blue) in the cells described in **C**. **(F)** Representative images of U2OS control and DFCP1 KO cells expressing the lipase-dead mutant (D166G) of ATGL (BFP-ATGL^DG^). Cells were treated with OA and starved as in **A**. **(G)** Colocalization (Pearson’s correlation coefficient, *rp*) of GFP-ATGL^DG^ with LDs in cells described in **F**. **(H)** Representative images of U2OS control and DFCP1 KO cells expressing BFP-ATGL^DG^ and either GFP or GFP-DFCP1^WT^. Cells were treated with OA and starved as in **A**. (I) Colocalization (Pearson’s correlation coefficient, *rp*) of BFP-ATGL^DG^ with LDs in cells described in **H**. The scale bars in whole-cell and inset images represent 10 and 2 µm, respectively. The statistical significance of the measurements in **B, D, E, G, and I** was determined using the Mann–Whitney U-test on the indicated number of observations from two independent transfections. Exact *p*-values are reported with exception to *p*>0.05, which is considered to be significant (n.s.).

This DFCP1-dependent localization of ATGL to LDs could arise directly through the localization of DFCP1, or through a DFCP1-dependent change to the pool of LDs, which may make them more “available” to ATGL. To distinguish between these possibilities, we exploited the observation that the localization of DFCP1 to LDs depends on its interaction with nucleotides (8). In particular, we have previously shown that interfering with the ability of DFCP1 to bind nucleotide through introduction of a point mutation (K193A) in the NTPase domain of DFCP1 (DFCP1^KA^) also impaired the ability of DFCP1 to translocate from the ER on to LDs in Hep3B cells (8). This is also the case for U2OS cells, in the absence (**Supplemental Figure 1D and 1E**) and presence (**Supplemental Figure 1C**) of exogenous GFP-ATGL expression. Thus, if ATGL localization depends on DFCP1 localization, we would expect to see less recruitment of ATGL to LDs in cells expressing the K193A mutant. Indeed, overexpression of BFP-DFCP1^KA^ resulted in significantly lower colocalization of ATGL to LDs when compared to those cells expressing BFP-DFCP1^WT^ **(Figure 2C and 2D)**. Importantly, GFP-ATGL colocalized equally well to both BFP-DFCP1^WT^ and BFP-DFCP1^KA^ in starved cells (**Figure 2E**) despite showing less LD localization (**Figure 2D**), which suggests that ATGL is either recruited to sites of DFCP1 accumulation, regardless of its location (LDs or ER), or is sensitive to a nucleotide-dependent structural state of DFCP1. Interestingly, this recruitment effect only occurs during starvation since there was not a statistically significant difference in the recruitment of ATGL to LDs in the presence of either BFP-DFCP1^WT^ or BFP-DFCP1^KA^ under fed conditions **(Supplemental Figure 1F)**.

The localization of ATGL to LDs is also known to depend on the catalytic activity of ATGL and starvation-induced phosphorylation (20, 21). We therefore set out to determine if DFCP1-dependent localization of ATGL to LDs also depends on one or more of these factors. To test whether the lipolytic activity of ATGL is necessary for its recruitment to DFCP1 coated LDs, we introduced a point mutation D166G in the catalytic domain of ATGL (ATGL^DG^) that was shown to disrupt a catalytic dyad in the N-terminal patatin-like domain (22). This so-called “lipase-dead” ATGL has been previously reported to stall when attempting to hydrolyze TAGs, thereby leading to pronounced ATGL localization to LDs in Cos7 cells (23). Similarly, we found enriched localization of GFP-ATGL^DG^ to and around LDs in U2OS cells (**Figure 2F and Supplemental Figure 1G**). Aside from having a broader distribution of LD localization under fed conditions, the sensitivity of GFP-ATGL^DG^ to starvation and DFCP1 was like that of GFP-ATGL^WT^ (**Figure 2G**). The localization of BFP-ATGL^DG^ to LDs was also invariant with DFCP1 expression in fed cells, whereas starvation increased the localization of BFP-ATGL^DG^ to LDs in starved control cells, but not in starved KO cells (**Figure 2F and 2G**). Despite this, overexpression of GFP-DFCP1 promoted the accumulation of BFP-ATGL^DG^ (**Figure 2H and 2I**) and, thus, the starvation dependent localization of GFP-ATGL is not dependent on its lipolytic activity, but instead depends on DFCP1.

Phosphorylation of ATGL was also not affected by DFCP1. Phosphorylation of S406 on ATGL by either AMPK or PKA was shown to be important for ATGL to mobilize on to LDs (20, 21). Since these two kinases are well-known to be activated by energetic stress and DFCP1-dependent recruitment of ATGL is more pronounced in starved cells, we tested the extent by which DFCP1 could influence ATGL phosphorylation. As commercially available primary antibodies sensitive to S406 phosphorylation are only available for mouse ATGL, we probed for S406 phosphorylation in control and DFCP1 CRISPR KO 3T3-L1 cells. While we do observe a pronounced starvation-dependent increase in phosphorylation of mouse ATGL, as expected, there was no observable difference to the extent of this phosphorylation in control and KO cells (**Supplemental Figure 1H**). Altogether, these results are surprising given that the localization of ATGL to LDs has been historically associated with its lipase activity and phosphorylation, but our data indicates that ATGL localization can be influenced by DFCP1, irrespective of these additional factors.

### DFCP1 Anchors ATGL to the LD

The dynamics of ATGL with LDs has been previously shown to correlate with the cellular activity of ATGL, such that activated ATGL associates and dissociates on LDs more quickly than inactive ATGL (24). Since DFCP1 influences the localization of ATGL in cells, we therefore wondered if DFCP1 could also impact ATGL dynamics, and, consequently, its activity in cells. To address this question, we examined dynamic changes of GFP-ATGL on LDs using fluorescence recovery after photobleaching (FRAP), in fed and starved U2OS cells **(Figure 3).** We found that GFP-ATGL in fed (**Supplemental Figure 2A and Supplemental Movie 1**) and starved (**Figure 2A and Supplemental Movie 2**) control cells recovered with a similar biphasic rate and to a similar extent (19% vs 18%) 15 min after photobleaching (**Figure 3A, 3E, and 3G**). By contrast, GFP-ATGL also recovered in a biphasic manner, but to a greater extent in both fed (35%) and starved (51%) DFCP1 KO cells, with the recovery of ATGL being significantly more than starved KO cells at 15 min after photobleaching (**Figure 3B, 3E, 3G, and Supplemental Figure 2B; Supplemental Movies 1 and 2**). This increase in GFP-ATGL dynamics likely reflects an increase in ATGL activity in the absence of DFCP1, since a kinase-dead mutant of GFP-ATGL^DG^ showed little recovery to LDs in both starved control and DFCP1-KO cells (**Figure 3C, 3F, 3G, and Supplemental Figure 2C; Supplemental Movie 3**). This enhancement of ATGL recovery was specific to DFCP1, since rescuing DFCP1 KO cells with BFP-DFCP1 reduced the extent of recovery to that of control cells **(Figure 3D, 3F, and 3G; Supplemental Movie 4).** Rescuing DFCP1 KO cells with DFCP1^KA^ also reduced the extent of GFP-ATGL recovery, but not to the same extent as WT DFCP1 (**Figure 3F, 3G, and Supplemental Figure 2D; Supplemental Movie 4**). It should be noted that in these rescue experiments we only examined the recovery of GFP-ATGL on LDs that were also coated with DFCP1, and thus this analysis does not account for the impaired localization of DFCP1^KA^ with LDs (**Supplemental Figure 1E**).

**Figure 3:**
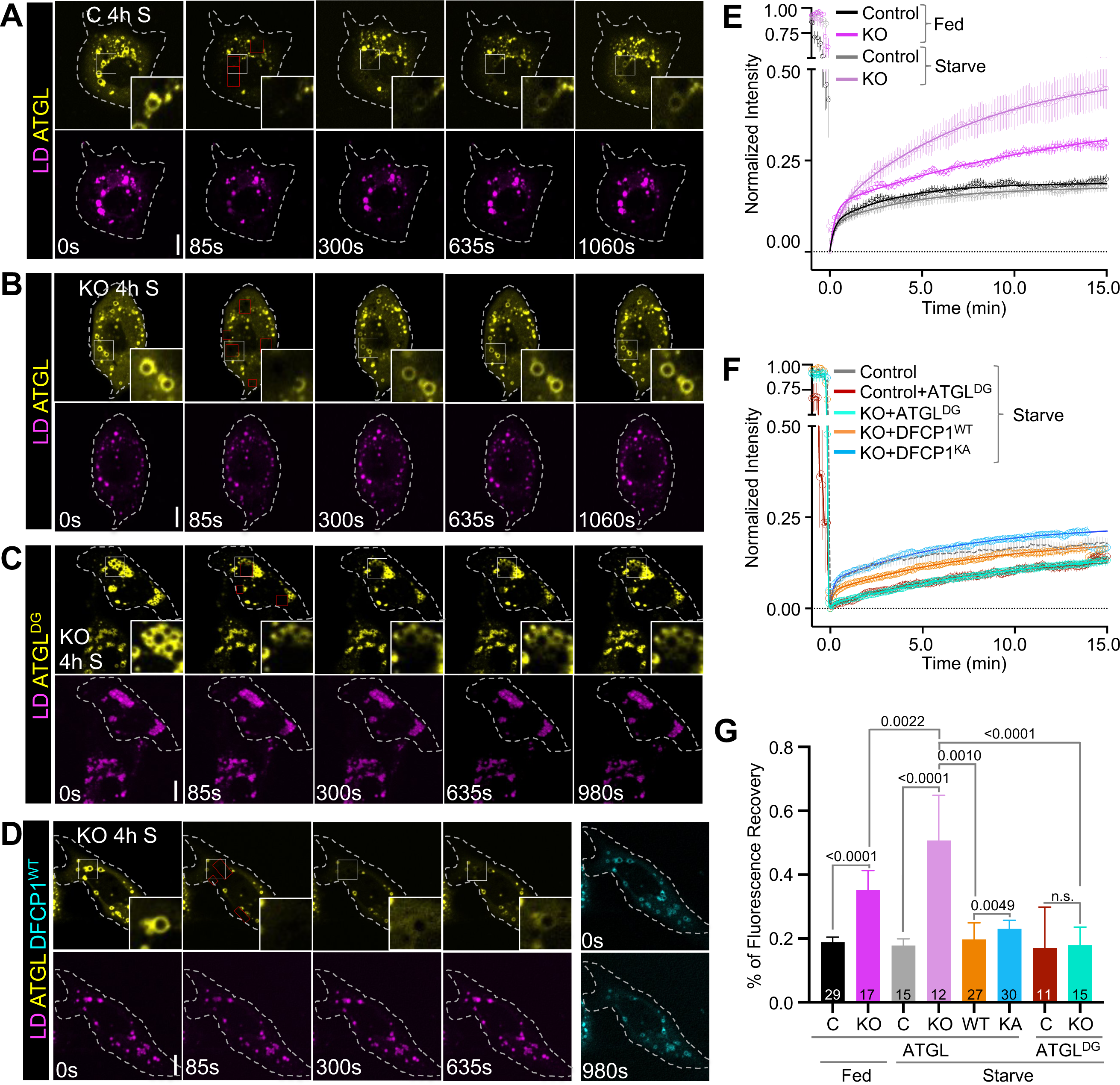
DFCP1 Anchors ATGL to the LD. **(A,B)** Fluorescence Recovery after Photobleaching (FRAP) of GFP-ATGL on individual LDs (shown in the inset) in control and DFCP1 KO U2OS cells that were stimulated with 200 µM OA for 20 h and starved (EBSS) for 4 h, with a subsequent treatment of LipidTOX Deep Red for 30 min. **(C)** FRAP of GFP-ATGL^DG^ on individual LDs (shown in the inset) in DFCP1 KO U2OS cells that were treated as in **A**. **(D)** FRAP of GFP-ATGL on individual LDs (shown in the inset) in DFCP1 KO U2OS cells rescued with BFP-DFCP1 and treated as in **A**. **(E)** Normalized average fluorescence recovery traces of GFP-ATGL in fed control cells (black), starved control cells (gray), fed DFCP1 KO cells (magenta), and starved DFCP1 KO cells (light magenta). Data is presented as mean ±SEM for the indicated number of traces (reported in **G**) and overlayed with a least-squares 2-component fit of the data (see methods). **(F)** Normalized average fluorescence recovery traces of GFP-ATGL^DG^ in starved control (brick red) and DFCP1 KO cells (cyan) and GFP-ATGL in starved DFCP1 KO cells rescued with BFP-DFCP1^WT^ (orange) or BFP-DFCP1^KA^ (blue). Data is presented and fitted as in **E**. **(G)** Extent of fluorescence recovery of GFP-ATGL^WT^ and GFP-ATGL^DG^ on LDs 15 min after photobleaching in fed (black) and starved (gray) control cells, fed (magenta) and starved (light magenta) KO cells, KO cells rescued with BFP-DFCP1^WT^ (orange) or BFP-DFCP1^KA^ (blue), and control (brick red) or KO (cyan) cells expressing GFP-ATGL^DG^. All scale bars in full cell and inset images represent 10 and 2 µm, respectively. The statistical significance of the measurements in **G** was determined using a two-tailed student t-test based on the indicated number of observations. Exact *p*-values are reported with exception to *p*>0.05, which is not considered to be significant (n.s.).

As suggested by the fluorescence recovery at 15 minutes, the rate of association for ATGL to LDs is markedly faster in DFCP1 KO cells. The average recovery trace for all conditions with GFP-ATGL fits well to a two-component association model (see methods). The first association rate was rapid and has a similar *t1/2* (∼0.25 min) across all conditions, which accounts for a small fraction (0.5-30%) of the total recovery. This fast recovery phase likely represents the fluorescence recovery of a cytosolic fraction of GFP-ATGL (**Supplemental Table 1**). By contrast, the second association rate was slow (**Supplemental Figure 2E**), and likely represents recovery of GFP-ATGL to the surface of LDs. However, the large variability in the *t1/2* values suggests that mobile fraction is not well defined at 15 min. Indeed, photobleached LDs were difficult to track beyond the 15-20 minutes and thus it is unlikely we were able to experimentally determine the true final fluorescence recovery value. Therefore, by assuming that GFP-ATGL can recover to at least the same extent as that seen in starved DFCP1 KO cells (51%), we found that the rate of GFP-ATGL was slow for control and DFCP1 rescue cells and was similar to the recovery of the enzymatically dead GFP-ATGL^DG^. By contrast, GFP-ATGL recovery was markedly faster for fed (*t1/2* = 16.2 ± 2.0 min) and starved (*t1/2* = 5.4 ± 0.38 min) GFP-ATGL DFCP1 KO cells (**Supplemental Figure 2F**). Additionally, DFCP1 KO cells rescued with BFP-DFCP1^KA^, showed a more rapid increase in GFP-ATGL than cells rescued with BFP-DFCP1 (*t1/2* = 27.7 ± 4.8 min vs. 33.2 ± 6.8 min). Taken together, this analysis suggests that DFCP1 functions as a tether that slows down the dynamic exchange of ATGL on the surface of LDs.

### DFCP1 Interacts with ATGL

Both the localization experiments (**Figure 2**) and FRAP experiments (**Figure 3**) suggest that DFCP1 can directly interact with ATGL. To test this possibility, we set out to pull down purified human ATGL using GFP-nanobody conjugated beads bound to GFP-DFCP1. Using this approach, we found that GFP-DFCP1 not only pulled down purified full-length ATGL **(Figure 4A and 4B),** but also an N-terminal truncation of ATGL (residues 1-254) that contains the patatin-like lipase domain (**Figure 4A** and **4C**). Interestingly, this region of ATGL plays a central role in its interactions with other ATGL interactors, such as CGI-58 (25). Using this more tractable fragment of ATGL, we determined that mutations in the NTPase domain of DFCP1 do not appear to affect the interaction with ATGL since DFCP1^KA^ pulls down with ATGL to the same extent as DFCP1^WT^ (**Figure 4C**), which is in strong agreement with our colocalization experiments (**Figure 2C and 2E**). Additionally, we found that a second mutation (R266Q) in the NTPase domain of DFCP1 (**Figure 4A**), which drives accumulation of DFCP1 on LDs (8), was also able to pull down ATGL to the same extent as DFCP1 (**Figure 4C**). Since these NTPase domain mutations do not influence binding of DFCP1 to ATGL, we conclude that the NTPase domain is not necessary for the physical interaction between ATGL and DFCP1 but remains important for the localization of DFCP1 to LDs. Indeed, by performing a systematic truncation analysis, we found that the N-terminal 415 residues of DFCP1 that include the NTPase domain, DFCP1^1-415^ (**Figure 4A**), was unable to pull down ATGL (**Figure 4D**). By contrast, truncations that contain the C-terminal FYVE domains, DFCP1^415-777^ or DFCP1^554-777^ (**Figure 4A**), were able to pull down ATGL (**Figure 4D**). Thus, the interaction between ATGL and DFCP1 requires the N-terminal lipase domain of ATGL and the C-terminal FYVE domains of DFCP1.

**Figure 4:**
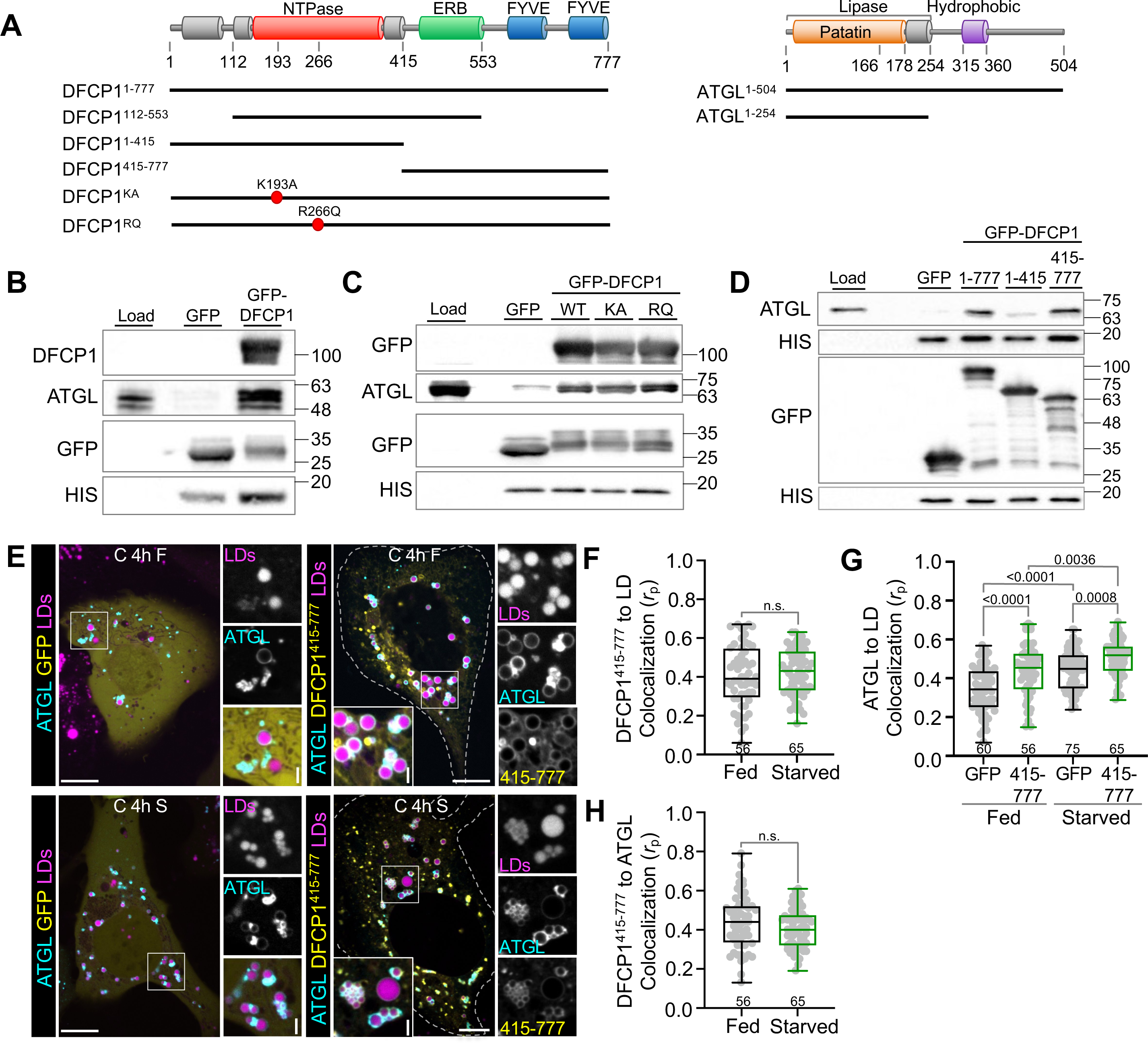
DFCP1 Interacts with ATGL. **(A)** Domain diagrams of DFCP1 and ATGL. DFCP1 contains an N-terminal Zinc-finger-like domain, an NTPase domain, and ER-binding (ERB) domain that binds to both the ER and LDs, and a pair of C-terminal PI3P-binding FYVE domains (FYVE). ATGL contains an N-terminal patatin-like lipase domain and a long-disordered region containing an LD-association domain. Constructs used in these studies are shown below each domain diagram. **(B)** CoIP of FLAG-ATGL using either GFP or GFP-DFCP1 constructs as bait, expressed in HEK 293T cells and pulled down onto HIS-tagged GFP-nanobody beads. **(C)** Pulldown using GFP or GFP-DFCP1^WT^, GFP-DFCP1^KA^, or GFP-DFCP1^RQ^ bound to HIS-tagged GFP-nanobody beads to test interaction with purified MBP-ATGL (1-254). **(D)** Pulldown using GFP, GFP-DFCP1^1-777^, GFP-DFCP1^1-415^, or GFP-DFCP1^415-777^ bound to HIS-tagged GFP-nanobody beads to test interaction with purified MBP-ATGL^1-254^. **(E)** Representative images of U2OS cells expressing eithr GFP or GFP-DFCP1^415-777^ and BFP-ATGL, treated with 200 µM OA for 20 h and then fed (basal growth media) or starved (EBSS) for 4 h, with a subsequent treatment of LipidTOX Deep Red for 30 min. **(F)** Colocalization (Pearson’s correlation coefficient, *rp*) of BFP-ATGL with LDs in fed (black) or starved (green) cells expressing GFP-DFCP1^415-777^ and treated as in **E**. **(G)** Colocalization (Pearson’s correlation coefficient, *rp*) of GFP (black) or GFP-DFCP1^415-^777 (green) with LDs in fed and starved cells expressing BFP-ATGL and treated as in **E**. **(H)** Colocalization (Pearson’s correlation coefficient, *rp*) of GFP-DFCP1^415-777^ (green) with BFP-ATGL in fed (black) or starved (green) cells and treated as in **E**. The scale bars in whole-cell and inset images represent 10 and 2 µm, respectively. The statistical significance of the measurements in **F-H** was determined using the Mann– Whitney U-test on the indicated number of observations from three independent transfections. Exact *p*-values are reported with exception to *p*>0.05, which is not considered to be significant (n.s.).

To validate this interaction, we set out to determine if the colocalization of ATGL with LDs was influenced by the minimal LD-interacting fragment DFCP1^415-777^. This truncation localized well to LDs on its own but this localization was insensitive to starvation (**Figure 4E and 4F**), which is likely due to the absence of the NTPase domain (8). The coexpression of GFP-DFCP1^415-777^ with BFP-ATGL led to an increase in LD localization of ATGL in both fed and starved cells when compared to the expression of GFP alone (**Figure 4G**) Interestingly, despite GFP-DFCP1^415-777^ not showing a significant increase in LD localization upon starvation (**Figure 4F**), or change in its colocalization with ATGL (**Figure 4H**), there was a marked increase in the BFP-ATGL accumulation on LDs in starved cells when compared to fed cells. This suggests that this DFCP1^415-777^ may synergize with ATGL and/or other ATGL activating factors to promote the accumulation of ATGL on LDs.

### DFCP1 Inhibits ATGL-Dependent Hydrolysis of TAGs

We have shown that DFCP1 drives the localization and dynamics of ATGL. Whereas this is likely part of the mechanism that DFCP1 uses to regulate LD size, we cannot exclude that DFCP1 can also be a negative regulator of ATGL activity. To determine if DFCP1 impairs lipolysis by ATGL, we used thin-layer chromatography (TLC) to compare the time-dependent increase of TAGs in starved control and DFCP1 KO cells stimulated to form LDs with a fluorescent lipid, Bodipy-C12 (**Figure 5A**). When compared to starved control cells, starved KO cells showed a reduced time-dependent accumulation of TAGs, which is consistent with an increase in lipolysis to counteract the storage of TAGs. To further support this assertion, we examined the lipidomic profiles of LDs purified from OA-stimulated control and KO cells **(Figure 5B, 5C, and Supplemental Figure 3A, 3B)**. Under starved conditions, there was a considerable increase in the total amount of all observed DAG species relative to the total amount of all observed TAG species in KO cells, when compared to control cells (**Figure 5B**). There was also a similar pattern of enrichment and de-enrichment of specific TAGs and DAGs in both fed and starved KO cells, that was distinct from fed and starved control cells (**Figure 5C and Supplemental Figure 3A**). In particular, the most abundant DAG (18:1/18:1) - formed from the hydrolysis of the most abundant TAG (18:1/18:1/18:1) that was formed by OA stimulation - was acutely enriched in both fed and starved KO cells, with starved KO cells showing the most enrichment of this species. Furthermore, we did not observe any changes in the abundances of cholesterol esters across all conditions (**Supplemental Figure 3B**), which further supports that DFCP1 specifically regulates ATGL and not neutral lipid storage in general.

**Figure 5:**
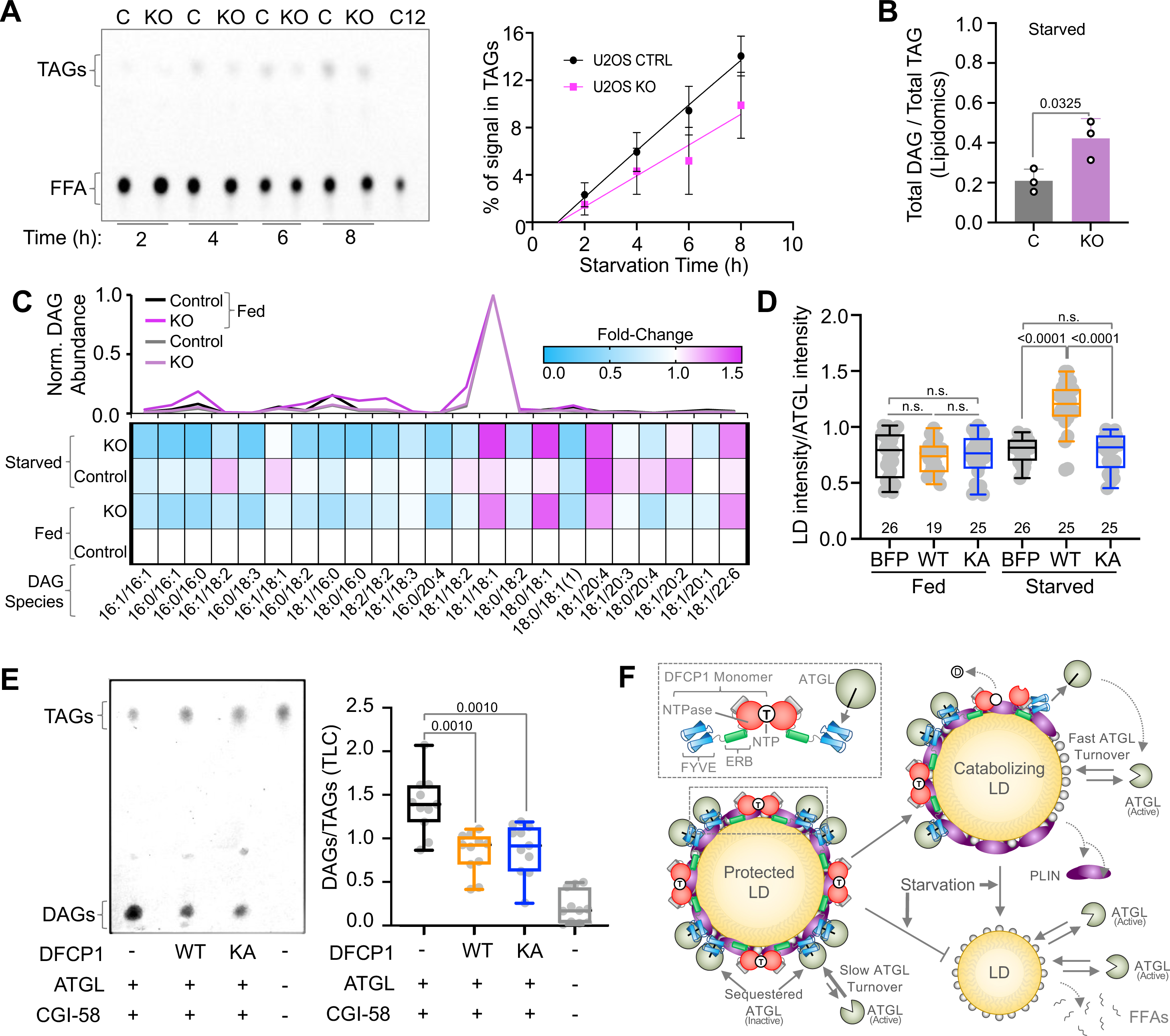
DFCP1 Inhibits ATGL-Dependent Hydrolysis of TAGs. **(A)** TLC plate showing the conversion of Bodipy C12 labeled TAGs into free Bodipy C12 FAs in control and DFCP1 KO U2OS cells that were treated with 2 µM Bodipy C12 in EBSS for the indicated time interval. The C12 lane shows free Bodipy C12 blotted on the TLC plate immediately before mobilization. Quantification of the accumulation of TAG levels in control and DFCP1 KO cells for 3 independent experiments is shown on the right. **(B)** Total DAGs normalized to the total TAGs from an average of 3 lipidomic profiles of LDs purified from U2OS control (C) and DFCP1 KO (KO) cell that were treated with 200 µM OA for 20 h and starved for 4 h. **(C)** Heat map showing the enrichment or loss of individual DAG species from LDs purified from U2OS control and DFCP1 KO cells in **B**, relative to those DAGs found in LDs isolated from WT fed cells. The plot above the heat map shows the average abundance of each DAG species relative to the most abundant DAG (18:1/18:1) species found in the LDs for each condition. **(D)** LD content in U2OS cells expressing GFP-ATGL and either BFP, BFP-DFCP1^WT^ or BFP-DFCP1^KA^ and treated with 200 µM OA for 20 h and then fed or starved for 4 h, with a subsequent treatment of LipidTOX Deep Red for 30 min. Sum projections of each Z-stack of entire cells were taken and total LD intensity in each cell was normalized to intensity of ATGL in the cell. **(E)** Reconstitution of LD lipolysis. LDs purified from DFCP1 KO U2OS cells were incubated with clarified HEK 293T lysates expressing GFP-CGI-58, mCherry-ATGL, and either BFP, BFP-DFCP1^WT^, or BFP-DFCP1^KA^ for 16 h and the resulting neutral lipid species were separated on TLC. Quantification of DAGs normalized to TAGs from TLC plates shown on the right. **(F)** Model of the mechanism by which DFCP1 regulates ATGL. Upon starvation, DFCP1 promotes the recruitment of ATGL to the LD surface through an interaction between the N-terminal catalytic domain of ATGL with the DFCP1^415-777^. This interaction immobilizes ATGL on the LD surface and represses its activity. Loss of DFCP1 from the LD surface, frees ATGL and allows it to rapidly dissociate and reassociate with LDs, to promote their catabolism. The statistical significance of the measurements in **B** and **E** was determined using two-tailed student t-test on three independent experiments for each condition in **B** and three independent experiments and three technical replicates for **E**. The statistical significance of the measurements in **D** was determined using the Mann–Whitney U-test on the indicated number of observations from three independent transfections. Exact *p*-values are reported with exception to *p*>0.05, which is not considered to be significant (n.s.).

Since loss of DFCP1 promotes lipolysis, we then investigated whether overexpression of DFCP1 could protect LDs from ATGL-mediated degradation in cells. For this purpose, we examined the LD content in fed and starved U2OS cells overexpressing GFP-ATGL, normalized to the ATGL intensity. We normalize to ATGL intensity in this case because its well appreciated that the amount of LDs is inversely correlated with the abundance of ATGL (26). We found that transient expression of either BFP-DFCP1^WT^ or BFP-DFCP^KA^ did not lead to a significant increase in LDs (based on the intensity of the LD marker LipidTOX) relative to GFP-ATGL expression levels (based on the intensity of GFP-fluorescence) **(Figure 5D)**. However, during starvation, BFP-DFCP1^WT^, but not BFP or BFP-DFCP1^KA^, promoted the accumulation of LDs relative to the abundance of GFP-ATGL (**Figure 5D)**. Thus, DFCP1^WT^ overexpression can help to protect LDs and thereby promote accumulation of LDs, even when ATGL is overexpressed.

In cells, the activity of ATGL is enhanced by the LD resident protein, CGI-58, and thus we wanted to more directly test if DFCP1 can regulate a reconstituted physiologically relevant lipolytic reaction. To do so, we incubated purified LDs from DFCP1-KO cells with a combination of clarified HEK293T lysates from cells transiently expressing CGI58 and ATGL with either BFP, BFP-DFCP1^WT^, or BFP-DFCP1^KA^. The resulting products were chloroform extracted and run on TLC **(Figure 5E and Supplemental Figure 3C).** It should be noted that purified LDs by themselves showed very little accumulation of DAGs, whereas the addition of lysates from cells overexpressing ATGL and CGI-58, showed a marked increase in 2,3 DAG, but not 1,3 DAG (**Supplemental Figure 3C**). Consistent with our previous observations (**Figure 5D**), cell lysates containing similar abundances of DFCP1^WT^ or DFCP1^KA^ **(Supplemental Figure 3D)** had significantly lower levels of these DAGs, when normalized to the abundance of TAGs, indicating that DFCP1 directly impairs CGI-58-activated ATGL **(Figure 5E).** However, unlike the cellular context (**Figure 5D**), DFCP1^KA^, impaired lipolysis to the same extent as DFCP1^WT^, which is more consistent with our pulldowns that show the interaction between DFCP1 and ATGL does not depend on the nucleotide state. In other words, inability for DFCP1^KA^ to stimulate LD accumulation in cells is likely due to its mistargeting to cellular compartments that are likely absent in the clarified lysates used in the assay. Altogether, this data presents strong evidence that DFCP1 is both a critical recruitment factor and repressor of ATGL-dependent hydrolysis of TAGs.

## Discussion

LDs serve as vital energy reservoirs that can be readily tapped to meet the energy demands of the cell. Therefore, it is imperative that the TAGs housed within LDs can be rapidly broken down into FFAs to fuel ATP synthesis by mitochondria. ATGL is the rate-limiting enzyme involved in the first step of TAG hydrolysis, however, little is known about the molecular factors that regulate this enzyme. In this study, we expanded upon the emerging functions of DFCP1 on LDs to show that it is a regulator of ATGL-mediated lipolysis (**Figure 5G**). In particular, we show that loss of DFCP1 does not impact the LD biogenesis pathway, but mainly impairs pathways associated with LD catabolism. **(Figure 1)**. We also found that that DFCP1 impairs lipolysis by recruiting (**Figure 2 and Supplemental Figure 1**) and anchoring ATGL (**Figure 3 and Supplemental Figure 2)** to the LD surface. This recruitment involves binding between the N-terminal patatin-like domain of ATGL and the C-terminal FYVE domains of DFCP1 **(Figure 4)**. Ultimately, the interaction of DFCP1 with ATGL impairs hydrolysis of TAGs stored in LDs by LD-bound ATGL. In line with this observation, loss of DFCP1 markedly enhances ATGL mediated breakdown of LDs (**Figure 5 and Supplemental Figure 3**). When taken together, our data suggest that DFCP1 serves to prime ATGL for rapid LD catabolism by a two-step mechanism that consists of specific recruitment of ATGL to LDs by DFCP1, followed by inhibition of ATGL activity by DFCP1.

The discovery that DFCP1 regulates LD metabolism by regulating the recruitment and function of ATGL is a departure from previous observations that linked DFCP1 to the biogenesis of LDs (9, 15). In that work, it was demonstrated that DFCP1 acts at the initial stages of LD biogenesis (9, 15), possibly through Seipin (15). However, our results suggest that DFCP1 is instead playing a role in LD catabolism by regulating both the recruitment and activity of ATGL. While these two roles of DFCP1 are not mutually exclusive, it has been shown that ATGL deficiency results in a compensatory loss of enzymes involved in *de novo* lipogenesis, including acetyl-CoA carboxylase (ACC1), fatty acid synthase (FAS), ATP-citrate lyase (ATP-CL) and acetyl-CoA synthetase (AceCS1) (27), which ultimately lead to impaired LD biogenesis. Additionally, depending on how ATGL is activated, DFCP1 could impact LD biogenesis under a specific context, such as during basal lipolysis. ATGL produces mostly *sn*-1,3 DAGs, but CGI-58 alters ATGL’s stereoselectivity to produce *sn*-1,3 and *sn*-2,3 DAGs. Since DGAT2 has a preference for sn-1,3 DAGs, this suggests that DGAT2 may re-esterify DAG to TAG during basal lipolysis to reduce cytotoxicity, whereas DGAT1 may be important for both basal and CGI-58-stimulated lipolysis (28). Thus, DFCP1 may play more of a selective role in LD biogenesis during basal lipolytic breakdown. In any case, regulation of ATGL by DFCP1 is central to both the changes in LD biogenesis and catabolism.

Our observation that DFCP1 regulates LD dynamics through sequestration and inhibition of ATGL further raises the question how this DFCP1-ATGL interaction is released. One possibility is that it may compete with CGI-58 for interaction with ATGL. CGI-58 is arguably the best studied regulatory protein of ATGL where it has been established to play a key role in enhancing both the rate and stereoselectivity of TAG hydrolysis (28). CGI-58 is known to be sequestered on the surface of LDs under basal conditions by perilipins (29, 30). Upon lipolytic stimulation, this repression is relieved through a phosphorylation dependent mechanism (31) and CGI-58 can bind to LD-bound ATGL (30, 32). Consequently, this interaction greatly accelerates the *in vivo* activity of ATGL (7). Thus, it is possible that in addition to regulating the recruitment of ATGL, DFCP1 may prevent or disrupt the CGI-58-ATGL interaction on LDs. Indeed, DFCP1 can still inhibit ATGL activity even in the presence of CGI-58 in vitro (**Figure 5E**). In this context, DFCP1 may mirror the function of perilipin 5 (Plin5/PLPN5) in cardiomyocytes, where Plin5 was shown to recruit ATGL to LDs (33) and sequester it from interacting with CGI-58 (32). Interestingly, Plin5 can interact with both CGI-58 and ATGL, but not both at the same time. Rather, Plin5, which can oligomerize, can partition both CGI-58 and ATGL on LDs and thereby prime lipolysis by concentrating these proteins on the LD surface (34). Similarly, DFCP1 can also oligomerize, and could help to partition both ATGL and CGI-58 on the LD surface, to ready them for rapid LD catalysis. Since Plin5 is mainly expressed in highly oxidative tissues, it is possible that DFCP1 is a functional replacement in less oxidative tissues.

Another possibility is that DFCP1 may release ATGL in a nucleotide dependent manner. We have previously shown that mutations in the NTPase domain of DFCP1 modifies the localization of DFCP1 to both LDs and autophagosomes (8). Indeed, a mutation that prevents the binding of nucleotide (DFCP1^KA^), lead to a loss of DFCP1 from LDs, whereas mutations that promote dimerization (DFCP1^RQ^) lead to stabilization of DFCP1 on the LD surface. Despite this, both mutant forms of DFCP1 can pull-down ATGL equally well. Additionally, if DFCP1^KA^ is on LDs, it can repress the dynamics of ATGL in cells (**Figure 3F, 3G, Supplemental Figure 2D and Supplemental Movies 4**) and activity of ATGL in vitro (**Figure 5E**). However, in cells, DFCP1^KA^ does not accumulate as well as DFCP1^WT^ (**Supplemental Figure 1E**) and as a consequence DFCP1^KA^ is less able to drive ATGL accumulation on LDs (**Figure 2D**) and protect LDs from degradation (**Figure 5D**). Thus, regulation of ATGL by DFCP1 depends mainly on whether DFCP1 accumulates on LDs. At this time, it is unclear what factors stimulate DFCP1 catalytic activity, but autophagy-induced changes may play a role (8).

ATGL, like DFCP1, also plays a role in autophagy. Specifically, ATGL was shown to increase autophagy by promoting SIRT1 activity (35, 36). Furthermore, ATGL increases the accumulation of LC3 on LDs and enhances autophagic flux in the liver, which implies that ATGL impacts lipophagy as well as lipolysis (37). ATGL has also been suggested to be an autophagy adaptor, due to its potential LC3-interacting region (LIR) motifs where mutation of this motif displaces ATGL from the LD and disrupts lipolysis (38). As DFCP1 KD exhibits a modest effect on lipophagy and has been shown to localize to sites of autophagosome biogenesis, even in the presence of LDs (8), it is possible that DFCP1 inhibits ATGL specifically at certain populations of LDs tethered to the ER to coordinate lipophagy and lipolysis, although further study will be required to investigate this possibility.

In summary, we have shown that DFCP1 is a negative regulator of ATGL-mediated lipolysis. Specifically, we show that DFCP1 recruits ATGL to LDs and subsequently sequesters ATGL on LDs. However, under conditions that promote translocation of DFCP1 away from LDs, ATGL is able to drive rapid lipolysis of LD. Thus, we speculate that DFCP1 functions as a nutrient-sensitive molecular switch that can promote LD growth under basal conditions but potentiates LD catabolism during starvation.

## Supporting Information

3 Supplemental Figures, 2 Supplemental Tables and 4 Supplemental Movies.

## Acknowledgments

This work was supported by grants from the National Institutes of Health (R01 GM136925) and the Elsa U Pardee Foundation (P20-03924). We would like to thank Dr. Philipp Scherer (University of Texas Southwestern Medical Center) for the mouse adipose tissue genomic library that was used to clone CGI-58. The GFP-nanobody construct was a kind gift from BM Collins.

## Materials and Methods

### Mammalian Cell Culture

U2OS, Hep3B and 3T3L1 (preadipocyte) cells were cultured at 37 °C with 5% CO2 and in growth media consisting of MEM GlutaMax or DMEM GlutaMax Supplemented with 10% fetal bovine serum (FBS) and antibiotic-antimycotic (Thermo Fisher Scientific, Waltham, MA), respectively. Two days prior to live-cell imaging experiments, 50,000 cells were seeded on 3.5 cm imaging dishes and grown to 40% confluency prior to transfection with the indicated plasmids using FuGENE HD (Promega, Madison, WI). In some cases, cells were also incubated for 2 days before imaging with a siRNA mixture, consisting of 5 μL of RNAiMax (ThermoFisher Scientific) and 2.5 pmol of the specified siRNA. Cells were induced to form LDs by Supplementaling the growth media with 200 μM oleic acid (OA) dissolved in ethanol for 4 or 24 h (as indicated), before exchanging the media with either basal growth media (fed) or EBSS (starved) for 1, 4, or 18 h (as indicated) prior to imaging. LDs were identified by incubating with the LD specific dye LipidTox Deep Red (ThermoFisher Scientific) at volume ratio of 1:10,000 for 30 min prior to imaging.

### Cloning

Mutations in the DFCP1 (K193A and R266Q) and ATGL (D166G) were introduced using the QuikChange mutagenesis kit (Agilent, Santa Clara, CA).

*FLAG-ATGL:* Human ATGL (Uniprot ID: Q96AD5) was cloned into a custom vector, where the DNA sequence encoding GFP in a pEGFP-C1 vector was replaced with a DNA sequence encoding a FLAG-TEV sequence using the NheI and BglII restriction sites.

*MBP-ATGL (1-254).* Residues 1-254 of human ATGL (Uniprot ID: Q96AD5) were inserted into the pMal-c2e bacterial expression vector using the EcoRI and Sal1 restriction enzyme cleavage sites.

*GFP-CGI-58.* CGI-58 (Uniprot ID: Q9DBL9) was PCR amplified from cDNA from WT mouse brown fat pads, provided kindly by Dr. Philipp Scherer, and cloned into a pEGFP-C1 vector using the SacI and SalI restriction sites.

### U2OS DFCP1 Knock-Out Cell Line

The U2OS DFCP1 Knock-out cell line was generated using Alt-R® CRISPR/Cas9 system (IDT, Coraville, IA), as previously described (8). In brief, a custom guide RNA (UAGCAGUGAUCGAUACGGAA) was annealed to tracer RNA, bound to purified *S. pyrogenes* Cas9 nuclease (IDT), and electroporated into U2OS cells using an Amaxa 4D-Nucleofector (Lonza Biosciences, Basel, Switzerland). Electroporated cells were then sorted into 96 well plates and checked for DFCP1 expression after 72 hours using western blotting. Clonal populations that were depleted of DFCP1 were further analyzed by next generation sequencing and monoclonal populations were selected based on the presence of two different but equivalent levels of frame-shifting deletions (to ensure both alleles of DFCP1 were edited).

### Live Cell Imaging and Image Analysis

All cells were imaged using either a Nikon Ti2 inverted microscope equipped with a 100× (1.4 NA) Plan-Apo oil immersion objective and a Yokogawa CSU-W1 spinning disk confocal attached to a Hamamatsu ORCA-FLASH4.0 CMOS camera. Cells in either DMEM FluoroBrite (ThermoFisher Scientific) medium Supplemented with 5% FBS or EBSS lacking phenol red (MilliporeSigma) were imaged at 37 °C and 5% CO2. Images stacks were captured at 16-bit 2048 x 2044 resolution with an axial spacing of 0.2 μm using the Nikon Elements Software package. All images were captured blindly and randomly, which involved imaging the nearest cell to a random set of x-y coordinates that contained actin fluorescence (depending on the experiment). Captured images were blinded again and image analysis was performed using the software Fiji (https://imagej.net/Fiji). Specifically, lipid droplet number and diameters were scored manually and 2D/3D colocalization analysis was performed using the Fiji Coloc2 analysis tool. Specifically, LD diameter was measured in the plane where a given LD’s diameter was the largest and lipid droplet density was determined by dividing the total number of LDs by the area of the cell’s basement membrane.

### Cell Fixation

U2OS cells expressing GFP-DFCP1^WT^ or GFP-DFCP1^KA^ were grown over no. 1.5 round glass coverslips (Electron Microscopy Sciences) and fixed in 4 % w/v paraformaldehyde (Electron Microscopy Sciences) in PBS for 15 min at RT, rinsed with PBS, and permeabilized in 0.1 % v/v Triton X-100 (Pierce Biotechnology) Following rinsing with PBS, 1:2000 LipidTOX was added along with 0.25 μg/ml of 4′,6-diamidino-2-phenylindole (DAPI; Molecular Probes) for 15 min, and added to cells for 30 min at RT. After rinsing with PBS, coverslips were mounted in a solution of ProLong Glass Antifade Mountant (P36980, Invitrogen), and allowed to cure for 24 h before imaging.

### FRAP

Imaging was performed on an Andor Dragonfly spinning disk confocal system using an iXon Lite EMCCD camera and a Nikon Eclipse Ti2 inverted microscope. Localized photobleaching was performed with Andor Mosaic 3 DMD array using a 445 nm laser at 145 nW power was scanned across the designated region at 0.9 ms/μm^2^. For imaging, 405, 488 and 637 nm solid state lasers were used along with BFP 480/20 nm, GFP 525/20 nm and Far Red 700/20 nm emission filters. Images were obtained with a Leica 63×, 1.4 NA oil immersion objective. Photobleaching regions of interest was performed approximately 1 min into imaging. Image acquisition was performed every 5 s. All imaging was performed at 37 °C and 5% CO2.

### FRAP Analysis

FRAP traces were normalized by setting the maximum pre-photobleach fluorescence intensity of GFP-ATGL within a given ROI to 1 and the minimum fluorescence intensity after photobleaching to 0. Individual traces of a given condition were aligned to the post-photobleached minimum and averaged. Average fluorescent recovery traces were fitted using a 2-component association model to account for the fast recovery of a small fraction of cytosolic ATGL surrounding the photobleached LD, as well as the recovery of GFP-ATGL to the LD.

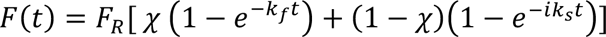

In this equation, *F*(*t*) is the normalized fluorescence intensity as a function of time, *FR* is the maximum recovered fluorescent intensity prior to photobleaching (normalized to 1 in this analysis), χ is the mol fraction, *kf* is the fast recovery rate, and *ks* is the slow recovery rate. All fits were performed using a weighted least squares fitting routine on the first 13 min of the data, and the mol fraction was allowed to vary but restricted to a maximum of 1; both the fast and slow recovery rates were allowed to vary. Additionally, in the comparisons of the GFP-ATGL recoveries at 15 min (**Supplemental Figure 2E**), both the recovery plateau and the slow rate were unrestricted and allowed to vary. In the comparison of the slow rates (**Supplemental Figure 2F**), the mobile fraction was fixed to the projected recovery of GFP-ATGL in DFCP1 KO cells for all traces, and only the slow rate and mol fraction were allowed to vary. Results of these fits are reported in **Supplemental Table 1**.

### Double Drug Experiment

Hep3B cells seeded on D35 imaging dishes treated with either non-targeting (NT) or DFCP1-targeting siRNAs (KD). After 48 h, control and DFCP1 KD cells were transfected with GFP using FuGENE and then, 4 h later, treated with 200 μM OA for 4 h. The OA was removed by washing cells twice with PBS before placing cells in starvation media (EBSS) Supplemented with either a combination of two LD metabolism drugs or an equivalent volume of DMSO. To study the impact of DFCP1 KD on LD biogenesis, both lipolysis and lipophagy were inhibited by treating starved cells with 20 μM ATGLi (Cayman #15284) and 50 μM Lalistat 2 (LALi; Cayman #25347). To study the impact of DFCP1 on lipolysis, both biogenesis and lipophagy were inhibited by treating starved cells with 10 μM DGAT1i (Sigma-Aldrich, PF-04620110) and 50 μM LALi. To study the effect of DFCP1 on lipophagy, both biogenesis and lipophagy were inhibited by treating starved cells with 10 μM DGAT1i and 20 μM ATGLi. Cells were treated with 1:10,000 LipidTOX Deep Red after 18 h incubation with the inhibitors and imaged.

### Cell-based TLC Lipase Assay

Cells were seeded on 10 cm dishes and then treated with a non-targeting siRNA or an siRNA for DFCP1 in complex with RNAimax. After 48 h, cells were washed with PBS and incubated with 3 μM BODIPY-C12 558/568 in EBSS for the indicated durations in **Figure 5A**. Cells were harvested via scraping and spun down in PBS. Cellular lipids were extracted using chloroform and spotted on aluminum-backed silica plates (Sigma) and then developed using 1:2 cyclohexane:ethyl acetate. Plates were imaged using BioRad Imager and spots were quantified using ImageJ.

### Phosphorylation Experiment

3T3L1 mouse preadipocytes were cultured to confluency in growth media (DMEM Glutamax with 10% FBS) and LD formation was stimulated by adding 200 µM OA to the media. After 20 h, media was removed, cells were washed with PBS, and cells were incubated in either fresh growth media (fed) or EBSS (starved) for 24 h. Prior to harvesting, cells were incubated with 0.1 µM calyculin A (LC Laboratories; C-3987) for 15 min at 37 °C to inhibit serine phosphatases. Cells were then harvested in PBS by scraping and lysed in lysis buffer containing 1% Triton-X, 1X PMSF, 0.1 µM calyculin A, 1X protease inhibitors (Pierce protease inhibitor tablets, EDTA free; A32965), 1X phosphatase inhibitors (Pierce phosphatase inhibitor tablets; A32957), 20 mM Tris, pH 7.5) and freeze-thawed once with liquid nitrogen.

### Western Blotting

Cells for western blotting were treated the same way as those for imaging, except, in this case, 200,000 cells were seeded onto a 6 cm plate one day prior to transfection with plasmids or siRNAs. Cells were harvested 3 days post-seeding in lysis buffer consisting of TBS Supplemented with 1% Triton-X, 2 mM EDTA with 1 mM PMSF. The lysis mixture was pipette mixed and incubated on ice for 30 min. The lysate was then clarified at 12,000 × *g* for 10 min and the supernatant was mixed with 3X SDS sample buffer. Samples were run on 12% polyacrylamide gels and transferred onto Immobilon-P polyvinylidene difluoride (PVDF) membranes (MilliporeSigma) membrane in transfer buffer containting 10% ethanol and 0.01% SDS at 130 mA for 3 h at 4 °C. Membranes were allowed to dry and then blocked for 1 h with 5% BSA in TBS Supplemented with 0.1% Tween (TBS-T), before incubating with the specified primary antibodies for 4 h to overnight at 4 °C. Blots were then washed with TBS-T and incubated with horseradish peroxidase (HRP)-linked anti-mouse and anti-rabbit immunoglobulin G secondary antibodies for 30 min at room temperature. Washed blots were developed using the clarity western ECL substrate (Bio-Rad Laboratories, Hercules, CA) and imaged using a Gel-Doc Imager (Bio-Rad). The relative abundance of proteins was determined by densitometry analysis using the program Fiji. The DFCP1 (85156S) and ATGL (2138S) antibodies were obtained from Cell Signaling Technologies (CST). The anti-phospho-ATGL (S406) antibody was obtained from Abcam (ab135093). The anti-CGI-58 antibody was obtained from ProteinTech (00041386). The anti-14-3-3 (133233) and anti-GFP (sc9996) antibodies were obtained from Santa Cruz Biotechnologies. The anti-HIS antibody (MA1 21315) was purchased from Invitrogen.

### Cell Viability Assays

U2OS cells were plated at an initial density of 300,000 cells/well in a 12-well plate, allowed to become 80-100% confluent, and then treated with 200 μM of OA or palmitic acid (PA) for 20 h followed by 4 h or 24 h of growth media (DMEM with 10% FBS) or EBSS. Cells were washed with 1 mL of PBS and then lifted with 300 μL trypsin-EDTA followed by quenching the trypsin with 600 μL of growth media. Cells were pelleted at 300 x *g* for 5 min and 600 μL of the supernatant was removed. The remaining 300 μL of solution was used to resuspend the cells. An equal volume of trypan blue was added, and living and dead cells were counted.

### Annexin V-FITC/PI Staining

For annexin V staining, cells were incubated in 90 μg/mL annexin V in annexin V binding buffer consisting of 10 mM HEPES, pH 7.4, 140 mM sodium chloride, 2.5 mM calcium chloride for 15 min at room temperature prior to recording fluorescence on a Citation 5 microplate reader (Agilent-BioTek, Santa Clara, CA), exciting at 488 nm and measuring emission at 535 nm. For propidium iodide staining, cells were incubated in 500 μg/mL propidium iodide in annexin V binding buffer consisting of 10 mM HEPES, pH 7.4, 140 mM sodium chloride, 2.5 mM calcium chloride for 15 mins at RT prior to examining fluorescence on a Citation 5 microplate reader (Agilent-BioTek, Santa Clara, CA), exciting at 535 nm and measuring emission at 610 nm. Measurements were normalized to DAPI fluorescence after fixation with methanol.

### *In vitro* TLC Lipase Assay

HEK 293T cells on 15 x cm^2^ dishes were transfected with 3.33 μg of each construct for a final 10 μg of total DNA using 25 μL polyethylenimine (PEI; Polysciences, Warrington, PA). Cells were lysed mechanically in a hypotonic solution (20 mM Tris-HCl (pH 7.0) 1 mM PMSF and protease inhibitor cocktail) 2 days post-transfection using a 28 gauge needle. HEK 293T lysates were clarified at 13,000 x *g* for 10 mins. Clarified lysates were normalized by total protein, determined by a Bradford assay, and then 500 μg of lysate was incubated with 10 μL of either control or DFCP1-KO LDs for the indicated length of time at 37 °C. After incubation, lipids were extracted with chloroform. The organic, solid, and aqueous phases were separated by spinning at 21,000 x *g* for 10 min at 4 °C and the organic phase was isolated. Separation of lipid species was achieved by spotting 1 μL of the solution on aluminum-backed silica plates (Sigma) and the plates were then developed using 70:30:1 hexane:diethyl ether:acetic acid. Visualization was done by brief incubation in 10% cupric sulfate in 8% aqueous phosphoric acid, followed by drying for 10 mins at RT and then heating on a hot plate at 160 °C. Plates were imaged using a LI-COR imager and spots were quantified using ImageJ.

### Protein Purification

#### FLAG-ATGL Purification

Human FLAG-ATGL (Uniprot ID: Q96AD5) was expressed in 20 15-cm dishes of HEK 293T cells were cultured in DMEM (ThermoFisher Scientific) with 5% FBS and then transiently transfected with a solution of 1 µg mL^-1^ plasmid DNA and 1 mg mL^−1^ PEI. After 48 h expression, cells were harvested in PBS, pelleted, and resuspended in lysis buffer comprised of 25 mM Tris-HCl (pH 7.0), 150 mM NaCl, 1% Triton-X, 1 mM PMSF, and a protease inhibitor cocktail (ThermoFisher Scientific). Cells were incubated in lysis buffer on ice for 30 min and insoluble cellular components were removed by centrifugation at 13,000 × *g* for 10 min. Clarified lysates were incubated with 1 mL Anti-DYKDDDDK (FLAG) Affinity Resin (Genscript; Piscataway, NJ) for 2 h at 4 °C and then washed with 10 column volumes (CVs) of wash buffer, comprised of 25 mM Tris, pH 7.0, 150 mM NaCl, and 1 mM PMSF, followed by 10 CVs of wash buffer Supplemented with 1 mM ATP. Bound ATGL was eluted by 4 sequential 1 h incubations of the FLAG resin with 1 CV of wash buffer Supplemented with 1 mg mL^-1^ FLAG peptide (prepared in house). The elution fractions were pooled, and the excess FLAG peptide was removed by dialysis using a 14 kDa MWCO dialysis tubing in a buffer composed of 20 mM Tris-HCl (pH 7.5), 150 mM NaCl and 1 mM DTT. The dialyzed protein was spin concentrated using a 30 kDa MWCO ultracentrifugation filter (MilliporeSigma) and subjected to size exclusion chromatography using a Superose 6 attached to an AKTA Pure 25L (Cytiva Life Sciences, Marlborough, MA) equilibrated in a buffer consisting of 20 mM Tris pH 7, 150 mM NaCl, and 1 mM DTT. The resulting peaks including ATGL were isolated together, spin concentrated using a 30 kDa MWCO ultracentrifugation filter, and flash frozen in liquid nitrogen.

#### MBP-ATGL (1-254) Purification

The MBP-ATGL (1-254) (Uniprot ID: Q96AD5) construct was transformed into *Rosetta* (DE3) cells (Novagen) that were subsequently cultured in Terrific Broth medium that was Supplemented with 100 µg mL^−1^ carbenicillin and 100 µg mL^−1^ chloramphenicol and incubated on a shaker at 37 °C. Protein expression was induced with 1 mM IPTG at 18 °C for 16 hr. Cells were homogenized in lysis buffer, comprised of 25 mM Tris pH 8.0, 300 mM NaCl, 4 mM benzamidine hydrochloride, and 1 mM PMSF, and lysed using a microfluidizer 110L (Microfluidics, Newton, MA). Lysates were clarified by centrifugation at 16,000 × *g* for 45 min and the supernatant was loaded onto amylose affinity resin (New England BioLabs, Ipswich, MA). The column was washed with 10 CVs of lysis buffer and the protein was eluted with sequentially added 1 CV aliquots of lysis buffer Supplemented with 20 mM maltose. Aliquots containing the eluted protein were combined and passed over a HiLoad 26/600 Superdex 200 pg size exclusion column (Cytiva) equilibrated in 20 mM Tris pH 8.0, 150 mM NaCl, and 1 mM DTT. The peak containing MBP-ATGL (1-254) was isolated, and the protein was spin concentrated to a 2 mL volume using a 10 MWCO ultracentrifugation filter. The protein was further purified with a MonoQ 4.6/100 PE column (Cytiva), using a 100 mM to 500 mM NaCl gradient spanning 30 CVs. The peak corresponding to MBP-ATGL (1-254) was isolated, spin concentrated, and then flash frozen in liquid nitrogen.

#### GFP Nanobody Purification

The GFP-nanobody construct was transformed into *BL21* (DE3) cells (Novagen) that were subsequently cultured in Terrific Broth medium that was Supplemented with 100 µg mL^−1^ carbenicillin and incubated on a shaker at 37 °C. Protein expression was induced with 1 mM IPTG at 18 °C for 16 h. Cells were homogenized in lysis buffer, comprised of 20 mM Tris pH 7.0, 300 mM NaCl, 4 mM benzamidine hydrochloride, and 1 mM PMSF, and lysed using a microfluidizer 110L (Microfluidics, Newton, MA) at 4 °C. Lysates were clarified by centrifugation at 12,000 × *g* for 45 min and the supernatant was loaded onto 1 mL of Nickel-NTA resin after adding imidazole to a final concentration of 10 mM. The Nickel-NTA resin was washed with 10 CV of lysis buffer Supplemented with 25 mM imidazole buffer. GFP-nanobody was eluted with lysis buffer Supplemented with 250 mM imidazole buffer and then dialyzed overnight into PBS and conjugated to 1 g of (N-hydroxysuccinimide) NHS-activated agarose beads (Pierce) overnight at 4 °C on a rocker. The reaction was quenched with 1M Tris, pH 8.0 for 15 min at RT. Conjugation efficiency was determined to be ∼95% and the concentration of GFP nanobody on the beads was determined to be 6 mg/mL.

### Lipidomics

Bligh-Dyer extraction was performed to extract TAG, PC, and CE from LD samples. The TAG (17:1/17:1/17:1), DAG (21:0/21:0/21:0), and d7-PE(18:2) were used as internal standards for TAG, DAG, and CE, respectively. Internal standard was added to the samples before extraction. Analysis of TAG, DAG, and CE were performed with a Shimadzu 20A HPLC system coupled to an API4000 mass spectrometer operated in positive multiple reaction monitoring (MRM) mode. Quality control (QC) samples were prepared by pooling aliquots of the study samples and were used to monitor the instrument performance. The QC samples were injected between every 4 study samples. Only the lipid species with coefficient of variance less than 15% in QC injections are reported. The relative quantification of lipids was provided, and the data were reported as the peak area ratios of the analytes to the corresponding internal standards (**Supplemental Table 2**). The relative quantification data generated in the same batch are appropriate to compare the change of an analyte in a test sample relative to other samples (*e.g.*, control vs. treated, or samples in a time-course study).

## References

1. Krahmer, N., Farese, R. V., and Walther, T. C. (2013) Balancing the fat: lipid droplets and human disease. EMBO Mol Med. 5, 973–983

2. Schulze, R. J., Sathyanarayan, A., and Mashek, D. G. (2017) Breaking fat: The regulation and mechanisms of lipophagy. Biochimica et Biophysica Acta (BBA) - Molecular and Cell Biology of Lipids. 1862, 1178–1187

3. Singh, R., and Cuervo, A. M. (2012) Lipophagy: Connecting Autophagy and Lipid Metabolism. International Journal of Cell Biology. 2012, 1–12

4. Schott, M. B., Weller, S. G., Schulze, R. J., Krueger, E. W., Drizyte-Miller, K., Casey, C. A., and McNiven, M. A. (2019) Lipid droplet size directs lipolysis and lipophagy catabolism in hepatocytes. Journal of Cell Biology. 218, 3320–3335

5. Kim, S.-J., Tang, T., Abbott, M., Viscarra, J. A., Wang, Y., and Sul, H. S. (2016) AMPK Phosphorylates Desnutrin/ATGL and Hormone-Sensitive Lipase To Regulate Lipolysis and Fatty Acid Oxidation within Adipose Tissue. Mol Cell Biol. 36, 1961–1976

6. Pagnon, J., Matzaris, M., Stark, R., Meex, R. C. R., Macaulay, S. L., Brown, W., O’Brien, P. E., Tiganis, T., and Watt, M. J. (2012) Identification and functional characterization of protein kinase A phosphorylation sites in the major lipolytic protein, adipose triglyceride lipase. Endocrinology. 153, 4278–4289

7. Lass, A., Zimmermann, R., Haemmerle, G., Riederer, M., Schoiswohl, G., Schweiger, M., Kienesberger, P., Strauss, J. G., Gorkiewicz, G., and Zechner, R. (2006) Adipose triglyceride lipase-mediated lipolysis of cellular fat stores is activated by CGI-58 and defective in Chanarin-Dorfman Syndrome. Cell Metabolism. 3, 309–319

8. Ismail, V. A., Naismith, T., and Kast, D. J. (2023) The NTPase activity of the double FYVE domain–containing protein 1 regulates lipid droplet metabolism. Journal of Biological Chemistry. 299, 102830

9. Li, D., Zhao, Y. G., Li, D., Zhao, H., Huang, J., Miao, G., Feng, D., Liu, P., Li, D., and Zhang, H. (2019) The ER-Localized Protein DFCP1 Modulates ER-Lipid Droplet Contact Formation. Cell Reports. 27, 343–358.e5

10. Gao, G., Sheng, Y., Yang, H., Chua, B. T., and Xu, L. (2019) DFCP1 associates with lipid droplets. Cell Biol Int. 43, 1492–1504

11. Xu, D., Li, Y., Wu, L., Li, Y., Zhao, D., Yu, J., Huang, T., Ferguson, C., Parton, R. G., Yang, H., and Li, P. (2018) Rab18 promotes lipid droplet (LD) growth by tethering the ER to LDs through SNARE and NRZ interactions. Journal of Cell Biology. 217, 975–995

12. Mei, S., Ni, H.-M., Manley, S., Bockus, A., Kassel, K. M., Luyendyk, J. P., Copple, B. L., and Ding, W.-X. (2011) Differential Roles of Unsaturated and Saturated Fatty Acids on Autophagy and Apoptosis in Hepatocytes. J Pharmacol Exp Ther. 339, 487–498

13. Tan, S. H., Shui, G., Zhou, J., Li, J. J., Bay, B.-H., Wenk, M. R., and Shen, H.-M. (2012) Induction of Autophagy by Palmitic Acid via Protein Kinase C-mediated Signaling Pathway Independent of mTOR (Mammalian Target of Rapamycin). Journal of Biological Chemistry. 287, 14364–14376

14. Axe, E. L., Walker, S. A., Manifava, M., Chandra, P., Roderick, H. L., Habermann, A., Griffiths, G., and Ktistakis, N. T. (2008) Autophagosome formation from membrane compartments enriched in phosphatidylinositol 3-phosphate and dynamically connected to the endoplasmic reticulum. Journal of Cell Biology. 182, 685–701

15. Lukmantara, I., Chen, F., Mak, H. Y., Zadoorian, A., Du, X., Xiao, F. N., Norris, D. M., Pandzic, E., Whan, R., Zhong, Q., and Yang, H. (2022) PI(3)P and DFCP1 regulate the biogenesis of lipid droplets. MBoC. 10.1091/mbc.E22-07-0279

16. Dow, R. L., Li, J.-C., Pence, M. P., Gibbs, E. M., LaPerle, J. L., Litchfield, J., Piotrowski, D. W., Munchhof, M. J., Manion, T. B., Zavadoski, W. J., Walker, G. S., McPherson, R. K., Tapley, S., Sugarman, E., Guzman-Perez, A., and DaSilva-Jardine, P. (2011) Discovery of PF-04620110, a Potent, Selective, and Orally Bioavailable Inhibitor of DGAT-1. ACS Med. Chem. Lett. 2, 407–412

17. Mayer, N., Schweiger, M., Romauch, M., Grabner, G. F., Eichmann, T. O., Fuchs, E., Ivkovic, J., Heier, C., Mrak, I., Lass, A., Höfler, G., Fledelius, C., Zechner, R., Zimmermann, R., and Breinbauer, R. (2013) Development of small-molecule inhibitors targeting adipose triglyceride lipase. Nat Chem Biol. 9, 785–787

18. Rosenbaum, A. I., Cosner, C. C., Mariani, C. J., Maxfield, F. R., Wiest, O., and Helquist, P. (2010) Thiadiazole Carbamates: Potent Inhibitors of Lysosomal Acid Lipase and Potential Niemann−Pick Type C Disease Therapeutics. J. Med. Chem. 53, 5281–5289

19. Bersuker, K., Peterson, C. W. H., To, M., Sahl, S. J., Savikhin, V., Grossman, E. A., Nomura, D. K., and Olzmann, J. A. (2018) A Proximity Labeling Strategy Provides Insights into the Composition and Dynamics of Lipid Droplet Proteomes. Developmental Cell. 44, 97–112.e7

20. Bartz, R., Zehmer, J. K., Zhu, M., Chen, Y., Serrero, G., Zhao, Y., and Liu, P. (2007) Dynamic Activity of Lipid Droplets: Protein Phosphorylation and GTP-Mediated Protein Translocation. J. Proteome Res. 6, 3256–3265

21. Ahmadian, M., Abbott, M. J., Tang, T., Hudak, C. S. S., Kim, Y., Bruss, M., Hellerstein, M. K., Lee, H.-Y., Samuel, V. T., Shulman, G. I., Wang, Y., Duncan, R. E., Kang, C., and Sul, H. S. (2011) Desnutrin/ATGL Is Regulated by AMPK and Is Required for a Brown Adipose Phenotype. Cell Metabolism. 13, 739–748

22. Duncan, R. E., Wang, Y., Ahmadian, M., Lu, J., Sarkadi-Nagy, E., and Sul, H. S. (2010) Characterization of desnutrin functional domains: critical residues for triacylglycerol hydrolysis in cultured cells. Journal of Lipid Research. 51, 309–317

23. Coassin, S., Schweiger, M., Kloss-Brandstätter, A., Lamina, C., Haun, M., Erhart, G., Paulweber, B., Rahman, Y., Olpin, S., Wolinski, H., Cornaciu, I., Zechner, R., Zimmermann, R., and Kronenberg, F. (2010) Investigation and Functional Characterization of Rare Genetic Variants in the Adipose Triglyceride Lipase in a Large Healthy Working Population. PLoS Genet. 6, e1001239

24. Schott, M. B., Rasineni, K., Weller, S. G., Schulze, R. J., Sletten, A. C., Casey, C. A., and McNiven, M. A. (2017) β-Adrenergic induction of lipolysis in hepatocytes is inhibited by ethanol exposure. Journal of Biological Chemistry. 292, 11815–11828

25. Cornaciu, I., Boeszoermenyi, A., Lindermuth, H., Nagy, H. M., Cerk, I. K., Ebner, C., Salzburger, B., Gruber, A., Schweiger, M., Zechner, R., Lass, A., Zimmermann, R., and Oberer, M. (2011) The Minimal Domain of Adipose Triglyceride Lipase (ATGL) Ranges until Leucine 254 and Can Be Activated and Inhibited by CGI-58 and G0S2, Respectively. PLoS ONE. 6, e26349

26. Smirnova, E., Goldberg, E. B., Makarova, K. S., Lin, L., Brown, W. J., and Jackson, C. L. (2006) ATGL has a key role in lipid droplet/adiposome degradation in mammalian cells. EMBO Reports. 7, 106–113

27. Lord, C. C., Ferguson, D., Thomas, G., Brown, A. L., Schugar, R. C., Burrows, A., Gromovsky, A. D., Betters, J., Neumann, C., Sacks, J., Marshall, S., Watts, R., Schweiger, M., Lee, R. G., Crooke, R. M., Graham, M. J., Lathia, J. D., Sakaguchi, T. F., Lehner, R., Haemmerle, G., Zechner, R., and Brown, J. M. (2016) Regulation of Hepatic Triacylglycerol Metabolism by CGI-58 Does Not Require ATGL Co-activation. Cell Reports. 16, 939–949

28. Eichmann, T. O., Kumari, M., Haas, J. T., Farese, R. V., Zimmermann, R., Lass, A., and Zechner, R. (2012) Studies on the substrate and stereo/regioselectivity of adipose triglyceride lipase, hormone-sensitive lipase, and diacylglycerol-O-acyltransferases. J Biol Chem. 287, 41446–41457

29. Shen, W.-J., Patel, S., Miyoshi, H., Greenberg, A. S., and Kraemer, F. B. (2009) Functional interaction of hormone-sensitive lipase and perilipin in lipolysis. Journal of Lipid Research. 50, 2306–2313

30. Granneman, J. G., Moore, H.-P. H., Granneman, R. L., Greenberg, A. S., Obin, M. S., and Zhu, Z. (2007) Analysis of Lipolytic Protein Trafficking and Interactions in Adipocytes. Journal of Biological Chemistry. 282, 5726–5735

31. Miyoshi, H., Perfield, J. W., Souza, S. C., Shen, W.-J., Zhang, H.-H., Stancheva, Z. S., Kraemer, F. B., Obin, M. S., and Greenberg, A. S. (2007) Control of Adipose Triglyceride Lipase Action by Serine 517 of Perilipin A Globally Regulates Protein Kinase A-stimulated Lipolysis in Adipocytes. Journal of Biological Chemistry. 282, 996–1002

32. Granneman, J. G., Moore, H.-P. H., Krishnamoorthy, R., and Rathod, M. (2009) Perilipin Controls Lipolysis by Regulating the Interactions of AB-hydrolase Containing 5 (Abhd5) and Adipose Triglyceride Lipase (Atgl). Journal of Biological Chemistry. 284, 34538–34544

33. Wang, H., Bell, M., Sreenevasan, U., Hu, H., Liu, J., Dalen, K., Londos, C., Yamaguchi, T., Rizzo, M. A., Coleman, R., Gong, D., Brasaemle, D., and Sztalryd, C. (2011) Unique Regulation of Adipose Triglyceride Lipase (ATGL) by Perilipin 5, a Lipid Droplet-associated Protein. Journal of Biological Chemistry. 286, 15707– 15715

34. Granneman, J. G., Moore, H.-P. H., Mottillo, E. P., Zhu, Z., and Zhou, L. (2011) Interactions of perilipin-5 (Plin5) with adipose triglyceride lipase. J Biol Chem. 286, 5126–5135

35. Khan, S. A., Sathyanarayan, A., Mashek, M. T., Ong, K. T., Wollaston-Hayden, E. E., and Mashek, D. G. (2015) ATGL-catalyzed lipolysis regulates SIRT1 to control PGC-1α/PPAR-α signaling. Diabetes. 64, 418–426

36. Lee, I. H., Cao, L., Mostoslavsky, R., Lombard, D. B., Liu, J., Bruns, N. E., Tsokos, M., Alt, F. W., and Finkel, T. (2008) A role for the NAD-dependent deacetylase Sirt1 in the regulation of autophagy. Proc. Natl. Acad. Sci. U.S.A. 105, 3374–3379

37. Sathyanarayan, A., Mashek, M. T., and Mashek, D. G. (2017) ATGL Promotes Autophagy/Lipophagy via SIRT1 to Control Hepatic Lipid Droplet Catabolism. Cell Reports. 19, 1–9

38. Martinez-Lopez, N., Garcia-Macia, M., Sahu, S., Athonvarangkul, D., Liebling, E., Merlo, P., Cecconi, F., Schwartz, G. J., and Singh, R. (2016) Autophagy in the CNS and Periphery Coordinate Lipophagy and Lipolysis in the Brown Adipose Tissue and Liver. Cell Metab. 23, 113–127

