## Supplemental Information for "DFCP1 is a Regulator of ATGL-mediated Lipid Droplet Lipolysis"

**Supplemental Table 1: FRAP Fit Result.**

**Supplemental Table 2: Lipidomic Profiles of Purified LDs**

**Supplemental Figure 1: DFCP1 Drives ATGL Localization in Cells**

**(A)** Cell viability measurements taken from U2OS cells treated with 200 μM oleic acid (OA, left) or palmitic acid (PA, right) for 20 h and then fed (basal growth media) or starved (EBSS) for the indicated amount of time.

**(B)** Propidium iodide (left) or Annexin V (right) fluorescence normalized to DAPI fluorescence on U2OS cells treated with indicated amount of OA or PA for 20 h and then fed or starved for 24 h.

**(C)** Colocalization (Pearson's correlation coefficient, $r_p$) of BFP-DFCP1$^{WT}$ (WT, orange) and BFP-DFCP1$^{KA}$ (KA, blue) with LDs in U2OS cells expressing GFP-ATGL and treated with 200 μM OA for 20 h before they were fed (basal growth media) or starved (EBSS) for 4 h and incubated with LipidTOX Deep Red for 30 min prior to imaging.

**(D)** Representative images of U2OS cells expressing either GFP-DFCP1$^{WT}$ or GFP-DFCP1$^{KA}$. Cells were treated with OA and starved as described in **C**.

**(E)** Colocalization (Pearson's correlation coefficient, $r_p$) of GFP-DFCP1$^{WT}$ (WT, black) and GFP-DFCP1$^{KA}$ (KA, blue) with LDs in cells described in **D**.

**(F)** Colocalization (Pearson's correlation coefficient, $r_p$) of GFP-ATGL with LDs, in fed KO cells rescued with either BFP (black), BFP-DFCP1$^{WT}$ (WT, orange) and BFP-DFCP1$^{KA}$ (KA, blue).

**(G)** Representative images of U2OS control and DFCP1 KO cells expressing BFP-ATGL$^{DG}$ and treated with 200 μM OA for 20 h before they were fed (basal growth media) and incubated of LipidTOX Deep Red for 30 min.

**(H)** Representative blot (left) of 3T3L1 mouse preadipocyte lysates extracted from control (black) and DFCP1 KO (magenta) cells treated with 200 μM OA for 20 h and then fed (basal growth media) or starved (EBSS) for 24 h. Densitometry of phosphorylated ATGL band relative to 14-3-3 band is shown on the right.

The scale bars in whole-cell and inset images represent 10 and 2 μm, respectively. Data from **A** are the result of at least three independent experiments with three technical replicates for each experiment. Data from **B** are from three technical replicates. Comparisons between control and DFCP1 KO cells in all conditions are not statistically significant as determined using a two-tailed student t-test on the indicated number of observations. The statistical significance of the measurements in **C, D,** and **E** was determined using the Mann–Whitney U-test on the indicated number of observations from two independent transfections. The statistical significance of the measurements in **F** was determined using a two-tailed student t-test on 5 replicates from two independent experiments. Exact *p*-values are reported with exception to $p$>0.05, which is not considered to be significant (n.s.).

**Supplemental Figure 2: DFCP1 Anchors ATGL to the LD.**
(**A,B**) Fluorescence Recovery after Photobleaching (FRAP) of GFP-ATGL on individual LDs (shown in the inset) in control and DFCP1 KO U2OS cells that were stimulated with 200 µM OA for 20 h and fed (basal growth media) for 4 h, with a subsequent treatment of LipidTOX Deep Red for 30 min.
**(C)** FRAP of kinase-dead mutant of ATGL (GFP-ATGL$^{DG}$) on a single LD (shown in the inset) in control U2OS cells that were stimulated with 200 µM OA for 20 h and starved (EBSS) for 4 h.
(**D**) FRAP of GFP-ATGL on a single LD (shown in the inset) in DFCP1 KO cells that were rescued with BFP-DFCP1$^{KA}$ and treated as in **C**.
(**E**) Time to half recovery ($t_{1/2}$) for the slow rate of fluorescence recovery for GFP-ATGL$^{WT}$ and GFP-ATGL$^{DG}$ on LDs in fed (black) and starved (gray) control cells, fed (magenta) and starved (light magenta) KO cells, KO cells rescued with BFP-DFCP1$^{WT}$ (orange) or BFP-DFCP1$^{KA}$ (blue), and control (brick red) or KO (cyan) cells expressing GFP-ATGL$^{DG}$. The slow rate was determined from least-squares 2-component "free" fit (see methods) of the data shown in Figures **3E** and **3F**. Data is presented as mean ±SD.
(**F**) Time to half recovery ($t_{1/2}$) for the slow rate of fluorescence recovery from least-squares 2-component fits (see methods) of the averaged data in Figures **3E** and **3F**, where the mobile fraction for all traces was assumed to be the same as that found in starved DFCP1 KO cells. Columns are labeled as indicated in **E** and the data is presented as mean ±SD for the indicated number of traces.

All scale bars in whole cell and inset images represent 10 and 2 µm, respectively. The statistical significance of the measurements in **E** and **F** was determined using the Mann–Whitney U-test. Exact *p*-values are reported with exception to $p$>0.05, which is not considered to be significant (n.s.).

**Supplemental Figure 3: DFCP1 Inhibits ATGL-Dependent Hydrolysis of TAGs.**
**(A)** Heat map showing the enrichment or loss of individual TAG species from LDs purified from U2OS control and DFCP1 KO cells in **Figure 1B**, relative to those TAGs found in LDs isolated from WT fed cells. The plot above the heat map shows the average abundance of each TAG species, relative to the most abundant TAG (18:1/18:1/18:1) species, found in the LDs harvested from each condition.
(**B**) Heat map showing the enrichment or loss of individual CE species from LDs purified from U2OS control and DFCP1 KO cells in **Figure 1B**, relative to those CEs found in LDs isolated from WT fed cells. The plot above the heat map shows the average abundance of each CE species, relative to the most abundant CE (16:1/18:2) species, found in the LDs harvested from each condition.
**(C)** TLC plate showing lipase activity after 16 h of incubation of LDs from DFCP1 KO U2OS cells, mixed with HEK 293T lysates expressing GFP-CGI-58, mCherry-ATGL, and either BFP-DFCP1$^{WT}$ (WT) or BFP-DFCP1$^{KA}$ (KA). LD lane shows the amount of DAGs

in LDs incubated with no HEK lysates. DAG lane is a control for the mobility of 1,3 DAGs and 2,3 DAGs.
**(D)** HEK 293T lysates showing the expression levels of GFP-CGI-58 (anti-GFP), mCherry-ATGL (anti-ATGL) , and BFP-DFCP1$^{WT}$ or BFP-DFCP1$^{KA}$ (anti-DFCP1).

**Supplemental Movie 1.** Representative FRAP Videos of GFP-ATGL in fed control (left) and DFCP1 KO (right) U2OS cells. Cells were treated as indicated in **Supplemental Figure 2A**. Photobleached regions are indicated by the white box.

**Supplemental Movie 2.** Representative FRAP Videos of GFP-ATGL in starved control (left) and KO (right) U2OS cells. Cells were treated as indicated in **Figure 3A**. Photobleached LDs are indicated by the white box.

**Supplemental Movie 3.** Representative FRAP Videos of kinase-dead mutant of ATGL (GFP-ATGL$^{DG}$) in starved control (left) and KO (right) U2OS cells. Cells were treated as indicated in **Figure 3C**. Photobleached LDs are indicated by the white box.

**Supplemental Movie 4.** Representative FRAP Videos of GFP-ATGL rescued with either BFP-DFCP1$^{WT}$ (left) or BFP-DFCP1$^{KA}$ (right) in starved control and DFCP1 KO U2OS cells. Cells were treated as indicated in **Figure 3D** and **Supplemental Figure 2D**. Photobleached LDs are indicated by the white box.

**Table S1: FRAP Fit Results**

Fit 1: 2-Component Association

| | % Recovery | | $\chi$ | | $k_f$ | | $k_s$ | | RMSE |
|---|---|---|---|---|---|---|---|---|---|
| | Mean | SE | Mean | SE | Mean | SE | Mean | SE | |
| Fed Control | 0.197 | 0.003 | 0.357 | 0.032 | 3.460 | 0.760 | 0.211 | 0.028 | 0.069 |
| Starve Control | 0.174 | 0.008 | 0.328 | 0.027 | 3.728 | 1.073 | 0.135 | 0.029 | 0.060 |
| Fed KO | 0.361 | 0.015 | 0.325 | 0.016 | 4.260 | 1.136 | 0.115 | 0.018 | 0.077 |
| Starve KO | 0.508 | 0.032 | 0.218 | 0.037 | 1.99 | 0.960 | 0.121 | 0.022 | 0.137 |
| Starve KO+DFCP1 | 0.197 | 0.010 | 0.246 | 0.013 | 6.565 | 2.427 | 0.114 | 0.016 | 0.054 |
| Starve KO+DFCP1$^{K1993A}$ | 0.230 | 0.005 | 0.332 | 0.009 | 6.962 | 1.220 | 0.142 | 0.011 | 0.041 |
| Starve Control+ATGL$^{D166G}$ | 0.191 | 0.021 | 0.065 | 0.010 | 3.15 | 0.671 | 0.072 | 0.014 | 0.037 |
| Starve KO+ATGL$^{D166G}$ | 0.187 | 0.015 | 0.080 | 0.010 | 6.59 | 6.021 | 0.074 | 0.011 | 0.035 |

Fit: 2-Component Association with Mobile set to 0.508

| | % Recovery | $\chi$ | | $k_f$ | | $k_s$ | | RMSE |
|---|---|---|---|---|---|---|---|---|
| | Mean | Mean | SE | Mean | SE | Mean | SE | |
| Fed Control | 0.508 | 0.236 | 0.005 | 2.460 | 0.149 | 0.018 | 0.001 | 0.064 |
| Starve Control | 0.508 | 0.208 | 0.006 | 2.460 | 0.149 | 0.017 | 0.001 | 0.064 |
| Fed KO | 0.508 | 0.203 | 0.007 | 2.460 | 0.149 | 0.045 | 0.001 | 0.064 |
| Starve KO | 0.508 | 0.208 | 0.010 | 2.460 | 0.149 | 0.125 | 0.003 | 0.064 |
| Starve KO+DFCP1 | 0.508 | 0.129 | 0.005 | 2.460 | 0.149 | 0.022 | 0.001 | 0.064 |
| Starve KO+DFCP1$^{K1993A}$ | 0.508 | 0.200 | 0.005 | 2.460 | 0.149 | 0.026 | 0.001 | 0.064 |
| Starve Control+ATGL$^{D166G}$ | 0.508 | 0.049 | 0.007 | 2.460 | 0.149 | 0.018 | 0.001 | 0.064 |
| Starve KO+ATGL$^{D166G}$ | 0.508 | 0.044 | 0.006 | 2.460 | 0.149 | 0.019 | 0.001 | 0.064 |

2-Component Association Fit Model: $F(t) = F_R[\, \chi\left(1 - e^{-k_f t}\right) + (1-\chi)(1 - e^{-ik_s t})]$

$\chi$: mol fraction

$k_f$: fast recovery rate

$k_s$: slow recovery rate

RMSE: Root Mean Square Error representing the goodness of fit

### Supplemental Figure 1

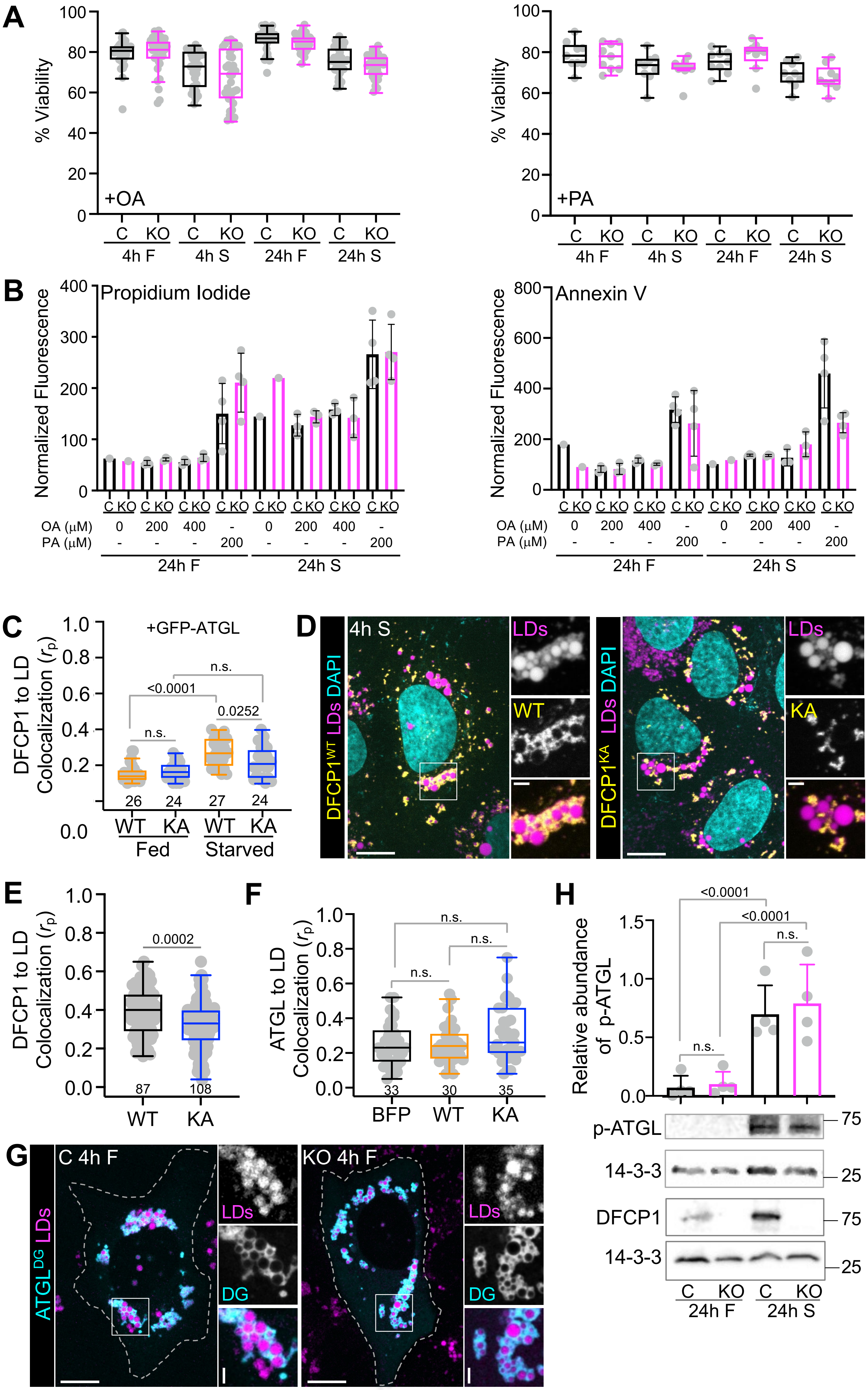

### Supplemental Figure 2

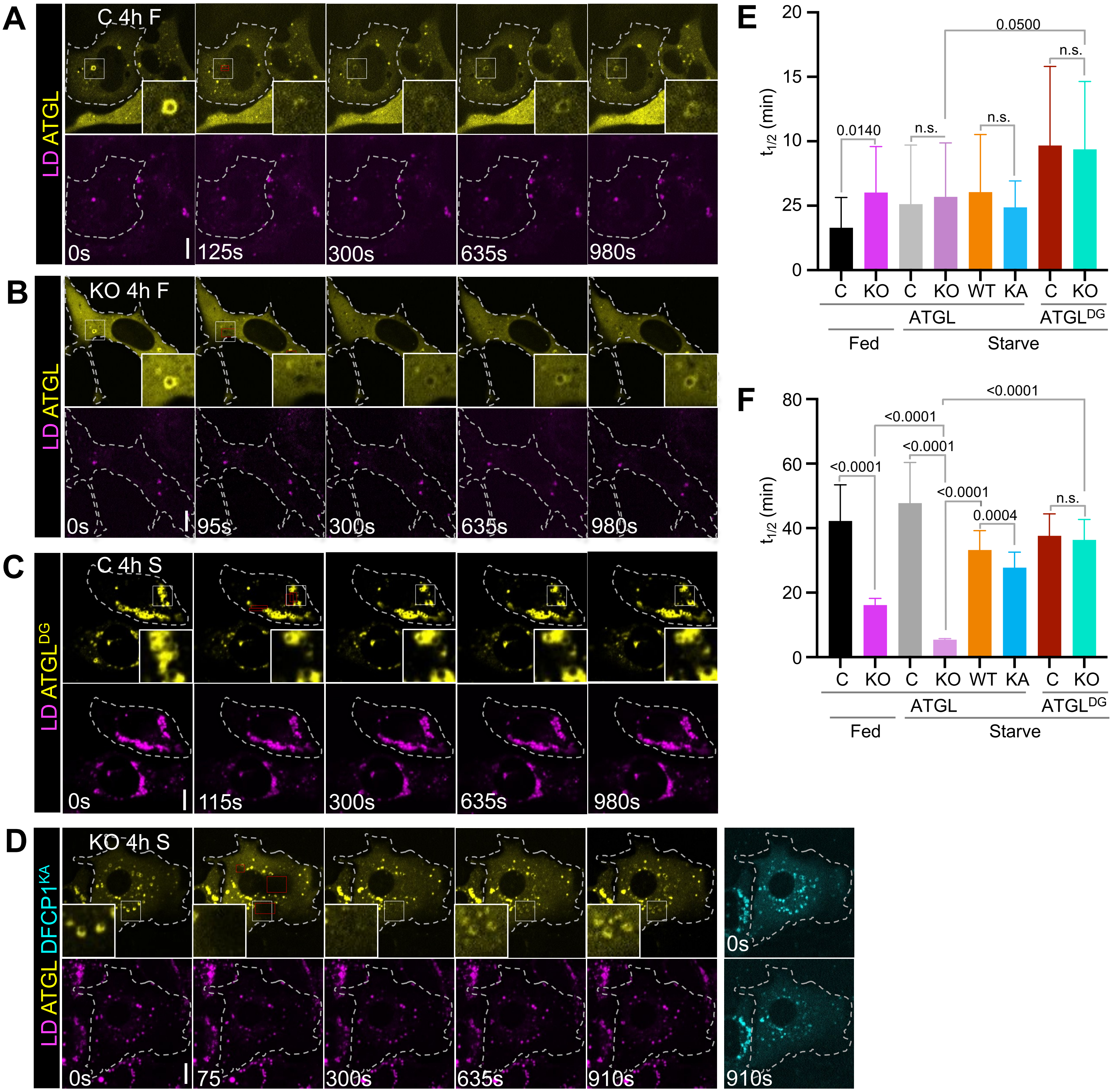

### Supplemental Figure 3

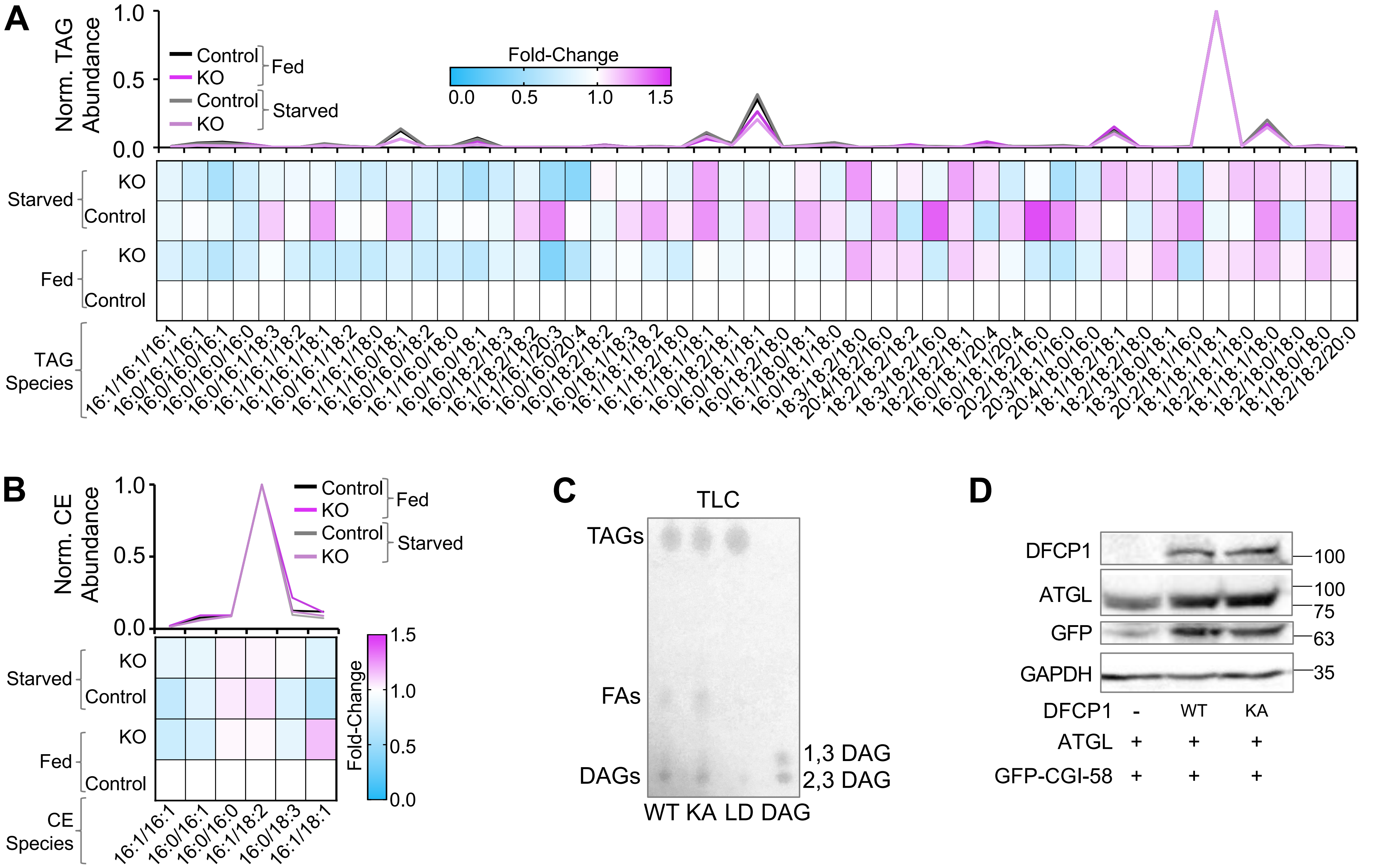
